# PseudotimeDE: inference of differential gene expression along cell pseudotime with well-calibrated p-values from single-cell RNA sequencing data

**DOI:** 10.1101/2020.11.17.387779

**Authors:** Dongyuan Song, Jingyi Jessica Li

## Abstract

To investigate molecular mechanisms underlying cell state changes, a crucial analysis is to identify differentially expressed (DE) genes along the pseudotime inferred from single-cell RNA-sequencing data. However, existing methods do not account for pseudotime inference uncertainty, and they have either ill-posed *p*-values or restrictive models. Here we propose PseudotimeDE, a DE gene identification method that adapts to various pseudotime inference methods, accounts for pseudotime inference uncertainty, and outputs well-calibrated *p*-values. Comprehensive simulations and real-data applications verify that PseudotimeDE outperforms existing methods in false discovery rate control and power.

## Introduction

In recent years, single-cell RNA-sequencing (scRNA-seq) technologies have undergone rapid development to dissect transcriptomic heterogeneity and to discover cell types or states in complex tissues [1, 2]. Embracing the capacity to measure transcriptomes of numerous cells simultaneously, scRNA-seq provides a powerful means to capture continuous cell-state transition across cells, and it has been used to study key cellular processes such as immune response [3] and cell development [4]. For example, a study of human fibroblasts identified distinct fibroblast subtypes responsible for mediating inflammation or tissue damage in arthritis [5]; a study of maternal-fetal interface tissue revealed new cell states and the importance of this tissue in maternal immune tolerance of paternal antigens [6]; a study of thymic development elucidated new principles of naïve T cell repertoire formation [7].

Pseudotime inference, also known as trajectory inference, is one of the most thriving scRNA-seq data analysis topics. The concept of “pseudotime” was first proposed in 2014 [8], and since then, more than 40 pseudotime inference methods have been developed [9]. Pseudotime inference aims to infer the ordering of cells along a lineage based on the cells’ gene expression profiles measured by scRNA-seq, and the inferential target is “pseudotime,” a time-like variable indicating the relative position a cell takes in a lineage. By establishing a temporal dimension in a static scRNA-seq dataset, pseudotime inference allows the probing of individual genes’ expression dynamics along with continuous cell-state changes. If a gene’s mean expression changes along pseudotime, the gene is referred to as differentially expressed (DE) and is likely to play an important role in the underlying cellular process that gives rise to the pseudotime. Identifying DE genes is the most crucial analysis after pseudotime inference because genes are the most fundamental functional units for understanding biological mechanisms.

Several methods have been developed to identify DE genes along inferred cell pseudotime. Popular pseudotime inference methods—TSCAN [10], Slingshot [11], Monocle [8], and Monocle2 [12]—include a built-in functionality for identifying DE genes after pseudotime inference. Their common approach is to use the generalized additive model (GAM) [13–15] to fit each gene’s expression level in a cell as a smooth-curve function of the cell’s inferred pseudotime. However, these built-in methods for DE gene identification are restricted as an add-on and downstream step of the pseudotime inference method in the same software package, and they cannot take external, user-provided pseudotime as input. Therefore, if users would like to use a new pseudotime inference method, they cannot use these built-in DE methods.

To our knowledge, only two DE gene identification methods can take any user-provided pseudotime. The first and state-of-the-art one is tradeSeq, which uses the negative binomial generalized additive model (NB-GAM) to model the relationship between each gene’s expression in a cell and the cell’s pseudotime [16]. Its *p*-value calculation is based on a chi-squared distribution, an inaccurate approximation to the null distribution. As a result, its *p*-values lack the correct probability interpretation. This issue is noted in the tradeSeq paper: “Rather than attaching strong probabilistic interpretations to the *p*-values (which, as in most RNA-seq applications, would involve a variety of hard-to-verify assumptions and would not necessarily add much value to the analysis), we view the *p*-values simply as useful numerical summaries for ranking the genes for further inspection.” Hence, the uncalibrated *p*-values of tradeSeq cannot be used for *p*-value-based statistical procedures such as the type I error control and the false discovery rate (FDR) control. The second method is Monocle3, better known as a pseudotime inference method [17], yet it also allows DE gene identification based on user-provided cell covariates via regression analysis. For clarity, we refer to the pseudotime inference and differential expression functionalities in Monocle3 as “Monocle3-PI” and “Monocle3-DE,” respectively. (Note that by “Monocle3-DE,” we mean the “regression analysis `fit_models()`,” not the “graph-autocorrelation analysis `graph_test()`,” in the Monocle3 R package; only the former works for user-provided pseudotime.) Monocle3-DE uses the generalized linear model (GLM) to identify DE genes for a user-provided covariate, e.g., pseudotime. However, GLM is more restrictive than GAM in that GLM assumes the logarithmic transformation of a gene’s expected read count in a cell is a strictly linear function of the cell’s pseudotime, while this assumption does not hold for many genes [18]. Hence, Monocle3-DE would miss those complex relationships between gene expression and pseudotime that do not satisfy its GLM assumption. In other words, Monocle3-DE’s restrictive GLM assumption impairs its power in identifying DE genes.

Besides the scRNA-seq methods we mentioned above, there are methods developed for identifying physical-time-varying DE genes from bulk RNA-seq time-course data. Among those methods, the ones allowing for continuous time can in principle be used to identify DE genes along pseudotime. Two examples of such methods are NBAMSeq [19] and ImpulseDE2 [20]. NBAMSeq is similar to tradeSeq in the use of NB-GAM, but it uses the Bayesian shrinkage method in `DESeq2` [21] to estimate gene variances, while tradeSeq does not. ImpulseDE2 [20], a method favorably rated in a benchmark study for bulk RNA-seq data [22], models gene differential expression by a unique “impulse” model. A later study modified ImpulseDE2 to identify DE genes along pseudotime from scRNA-seq data [16]. However, the performance of NBAMSeq and ImpulseDE2 on scRNA-seq data lacks benchmarking. Loosely related, many methods can identify DE genes between discrete cell clusters, groups, or conditions [23–27]; however, these methods are inapplicable to finding DE genes along continuous pseudotime.

More importantly, the existing methods that identify DE genes along pseudotime have a common limitation: they ignore the uncertainty of inferred cell pseudotime, which they consider as one fixed value per cell. This issue arises from the fact that most pseudotime inference methods only return point estimates of cell pseudotime without uncertainty quantification (i.e., every cell only receives an inferred pseudotime without a standard error), with few exceptions [28, 29]. Hence, downstream DE gene identification methods treat these point estimates as fixed and ignore their uncertainty. However, this ignorance of uncertainty would result in invalid *p*-values, leading to either failed FDR control or power loss. This critical problem has been noted in several pseudotime inference method papers [10, 11, 28] and in the tradeSeq paper [16], yet it remains an open challenge to our knowledge.

Motivated by the ill-posed *p*-value issue of existing pseudotime-based differential expression methods, we propose PseudotimeDE, the first method that accommodates user-provided pseudotime inference methods, takes into account the random nature of inferred pseudotime, and outputs well-calibrated *p*-values. PseudotimeDE uses subsampling to estimate pseudotime inference uncertainty and propagates the uncertainty to its statistical test for DE gene identification. As the most notable advantage of PseudotimeDE over existing methods, PseudotimeDE’s well-calibrated *p*-values ensures the reliability of FDR control and other downstream analyses, as well as avoiding unnecessary power loss due to overly-conservative *p*-values.

## Results

### Overview of the PseudotimeDE method

The statistical method of PseudotimeDE consists of four major steps: subsampling, pseudotime inference, model fitting, and hypothesis testing (Fig. 1). The first two steps are performed at the cell level and include all informative genes (whose selection depends on the pseudotime inference method, e.g., Slingshot and Monocle3-PI), while the last two steps are performed on every gene that is potentially DE.

1. In the subsampling step, PseudotimeDE subsamples 80% of cells from the original dataset to capture the uncertainty of pseudotime inference, the same technique as used in [9, 11, 30].
2. In the pseudotime inference step, PseudotimeDE applies a user-specified pseudotime inference method to the original dataset and each subsample, so that every cell receives its inferred pseudotime in the original dataset and all the subsamples that include it. To construct null cases where genes are non-DE for later hypothesis testing, PseudotimeDE permutes the inferred pseudotime in each subsample, independent of other subsamples.
3. In the model fitting step, PseudotimeDE fits NB-GAM or zero-inflated negative binomial GAM (ZINB-GAM) to every gene in the original dataset to obtain a test statistic that indicates the effect size of the inferred pseudotime on the gene’s expression.
4. In the hypothesis testing step, for every gene, Pseudotime fits the same model used for the original dataset to the permuted subsamples to obtain approximate null values of the gene’s test statistic (the null values are approximate because the subsamples do not have the same number of cells as in the original dataset). To save the number of subsamples needed and to improve the *p*-value resolution, Pseudotime fits a Gamma distribution or a mixture of two Gamma distributions to these null values. It subsequently uses the fitted parametric distribution as the approximate null distribution of the test statistic. Finally, PseudotimeDE calculates a right-tail *p*-value for the gene from the gene’s test statistic in the original dataset and the approximate null distribution.

**Figure 1:**
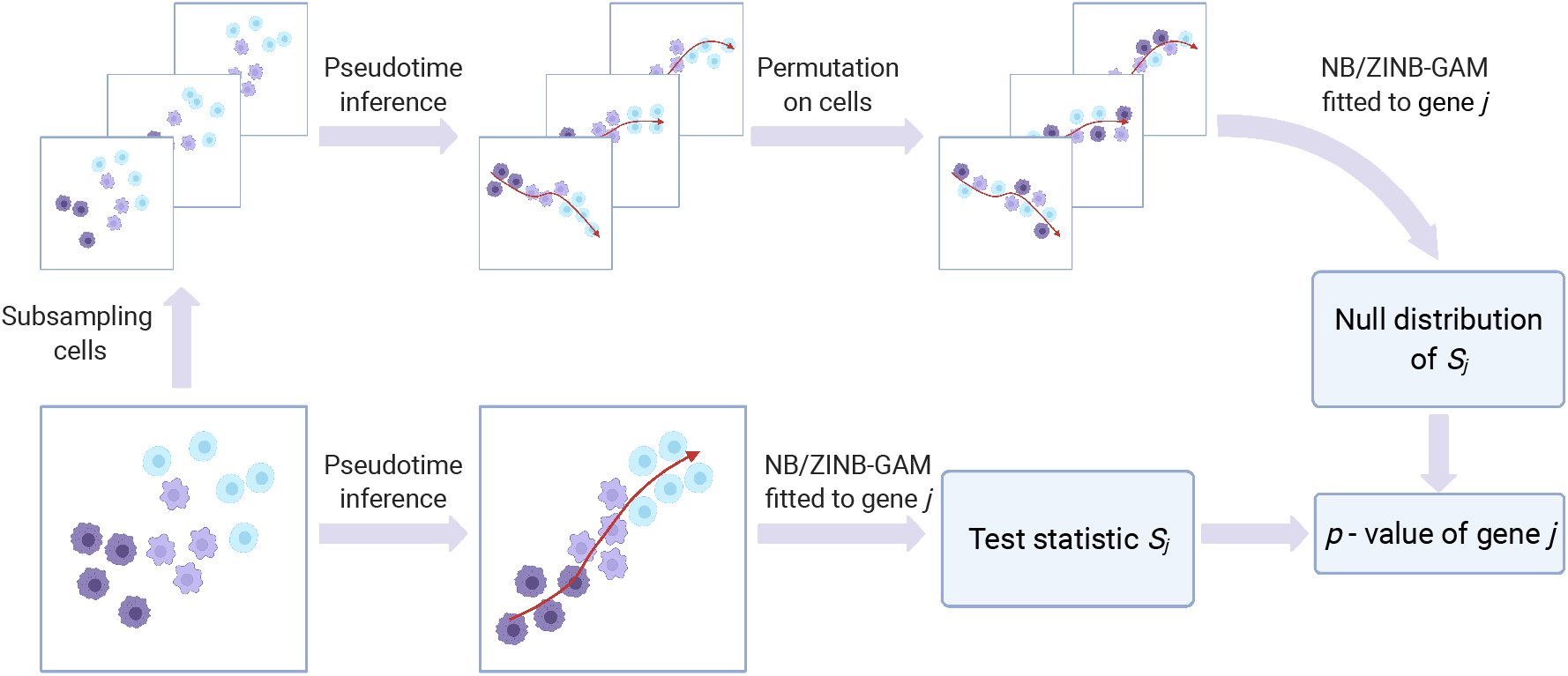
An illustration of the PseudotimeDE method. The core of PseudotimeDE is to obtain a valid null distribution of the DE gene test statistic *S_j_*. To achieve that, PseudotimeDE subsamples 80% cells from the original scRNA-seq data. Then on each subsample, PseudotimeDE performs pseudotime inference (using a user-specified method such as Slingshot and Monocle3-PI) and permutes the inferred pseudotime across cells. Next, PseudotimeDE fits a model (NB-GAM or ZINB-GAM) to the permuted subsamples to obtain the values of *S_j_* under the null hypothesis and uses these values to approximate the null distribution of *S_j_*. In parallel, PseudotimeDE fits the same model to the original dataset and calculate the observed value of *S_j_*. Finally, PseudotimeDE derives the *p*-value from the observed value and the null distribution of *S_j_*. Detail is described in Methods.

Further detail of PseudotimeDE is described in **Methods**.

### Simulations verify that pseudotimeDE outperforms existing methods in the validity of *p*-values and the identification power

We use a widely-used simulator `dyntoy` [9, 16] to generate four synthetic scRNA-seq datasets, among which three are single-lineage datasets with low-, medium- and high-dispersion levels, and the other is a bifurcation dataset. Since the single-lineage high-dispersion dataset best resembles the real scRNA-seq data (Additional file 1: Fig. S1), we use it as our primary case. We apply two pseudotime inference methods—Slingshot and Monocle3-PI—to each synthetic dataset to infer cell pseudotime.

First, we find that PseudotimeDE successfully captures the underlying uncertainty of inferred pseudotime. The first layer—”linear uncertainty”—reflects the randomness of inferred cell pseudotime within a cell lineage (Fig. 2a & c). Fig. 2b & d show the distributions of individual cells’ inferred pseudotime by Slingshot and Monocle3-PI, respectively, across 1000 subsampled datasets, confirming that linear uncertainty is specific to pseudotime inference methods. Between the two methods, Monocle3-PI demonstrates greater linear uncertainty. The second layer—”topology uncertainty”—reflects the randomness of lineage construction. The synthetic bifurcation dataset contains two cell lineages. Slingshot correctly constructs the bifurcation topology from the original dataset and the 1000 subsampled datasets. While Monocle3-PI captures the bifurcation topology from the original dataset (Fig. 2e), it fails to capture the topology from over 50% of subsamples (Fig. 2f shows randomly picked 10 subsamples), demonstrating its greater topology uncertainty than Slingshot’s.

**Figure 2:**
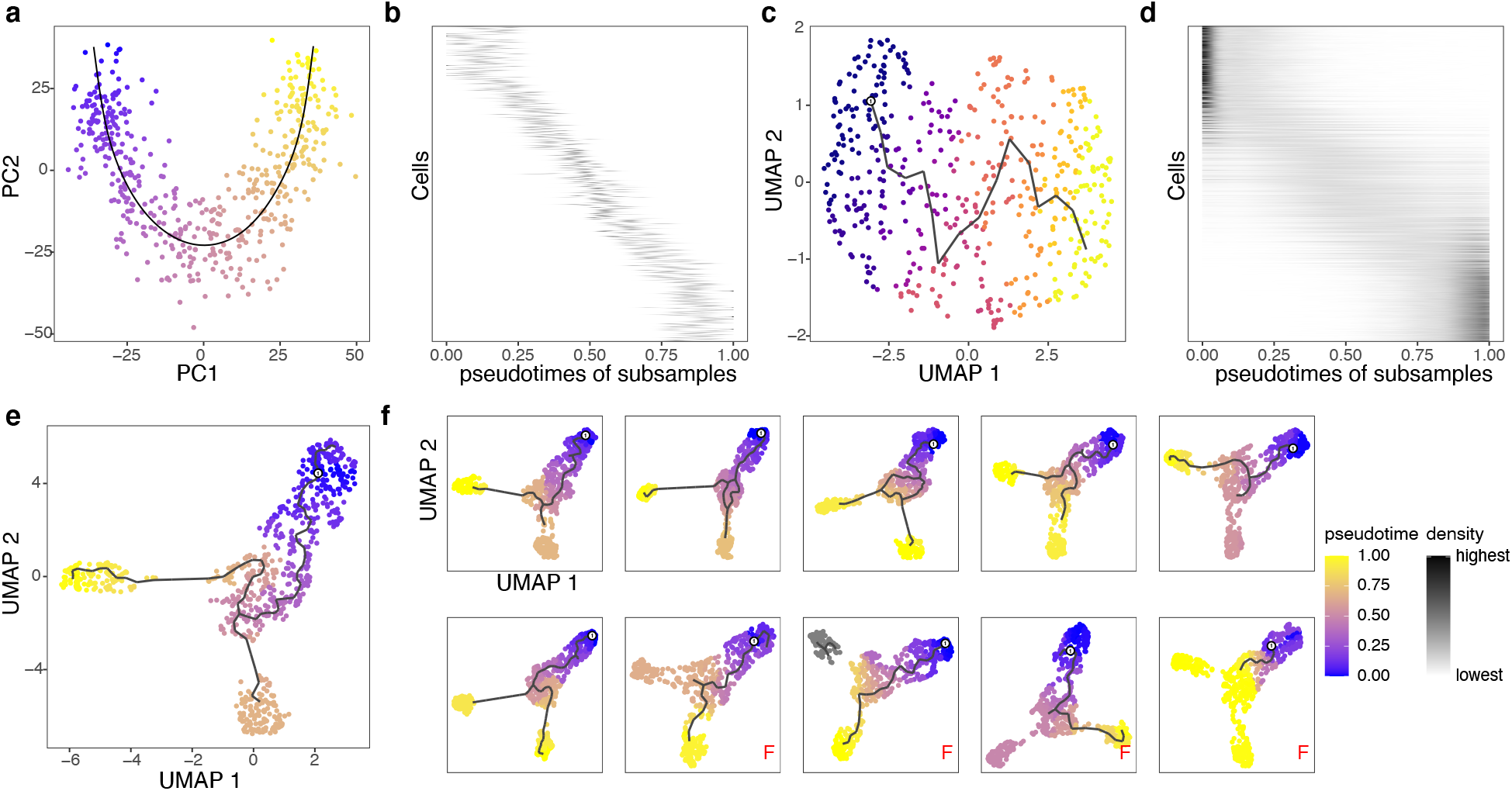
PseudotimeDE captures the uncertainty in pseudotime inference. **(a)** Visualization of synthetic singlelineage cells marked with inferred pseudotime by Slingshot (using PCA). The black curve denotes the inferred lineage. **(b)** The distributions of individual cells’ inferred pseudotime by Slingshot across subsamples. In the vertical axis, cells are ordered by their true time in the lineage used in simulation; for every cell (a vertical coordinate), black dots have horizontal coordinates corresponding to the cell’s inferred pseudotime in the subsamples that include the cell. The more horizontally spread out the black dots, the greater uncertainty the pseudotime inference has. **(c)** Visualization of synthetic single-lineage cells marked with inferred pseudotime by Monocle3-PI (using UMAP). The black curve denotes the inferred lineage. Compared with (a), the inferred lineage is more wiggling. **(d)** The distributions of individual cells’ inferred pseudotime by Monocle3-PI across subsamples. Compared with (b), the uncertainty in pseudotime inference is greater. **(e)** Visualization of synthetic bifurcating cells marked with inferred pseudotime by Monocle3-PI (using UMAP). Monocle3-PI recovers the bifurcation topology. **(f)** Visualization of ten subsamples of the cells in (e), marked with inferred pseudotime by Monocle3-PI (using UMAP) on each subsample. Four out of the ten subsamples do not have the bifurcation topology correctly inferred (labeled with red “F”), revealing the uncertainty in pseudotime inference by Monocle3-PI. In panels (a), (c), (e) and (f), inferred pseudotime is represented by a color scale from 0 (the earliest pseudotime) to 1 (the latest pseudotime).

After confirming pseudotime inference uncertainty, we benchmark PseudotimeDE against four DE gene identification methods: tradeSeq, Monocle3-DE, NBAMSeq, and ImpulseDE2. The first two methods, tradeSeq and Monocle3-DE, are the state-of-the-art for scRNA-seq data analysis and thus serve as the main competitors of PseudotimeDE. In our benchmark, we first evaluate these methods in terms of the validity of their *p*-values, which should be uniformly distributed between 0 and 1 under the null hypothesis (i.e., a gene is not DE). Our results show that, among the five methods, PseudotimeDE generates the best-calibrated *p*-values that follow the expected uniform distribution most closely (Fig. 3a & f and Additional file 1: Figs. S3–S5a & f). Among the existing four methods, only Monocle3-DE provides roughly calibrated *p*-values, while tradeSeq, NBAMSeq, and ImpulseDE2 output *p*-values that are much deviated from the expected uniform distribution. This observation is confirmed by the Kolmogorov–Smirnov test, which evaluates how closely *p*-values follow the uniform distribution. Since the identification of DE genes relies on a small *p*-value cutoff, the smaller *p*-values are more important than the larger ones. Hence, we re-plot the *p*-values on – log_10_ scale to closely examine the calibration of small *p*-values (Fig. 3b & g and Additional file 1: Figs. S3–S5b & g). Again, PseudotimeDE returns the best-calibrated *p*-values, while the other four methods generate overly small *p*-values that would inflate false discoveries. This is reflected in our results: at a target 5% FDR threshold, PseudotimeDE leads to the best FDR control among all methods (Fig. 3c & h and Additional file 1: Figs. S3–S5c & h).

**Figure 3:**
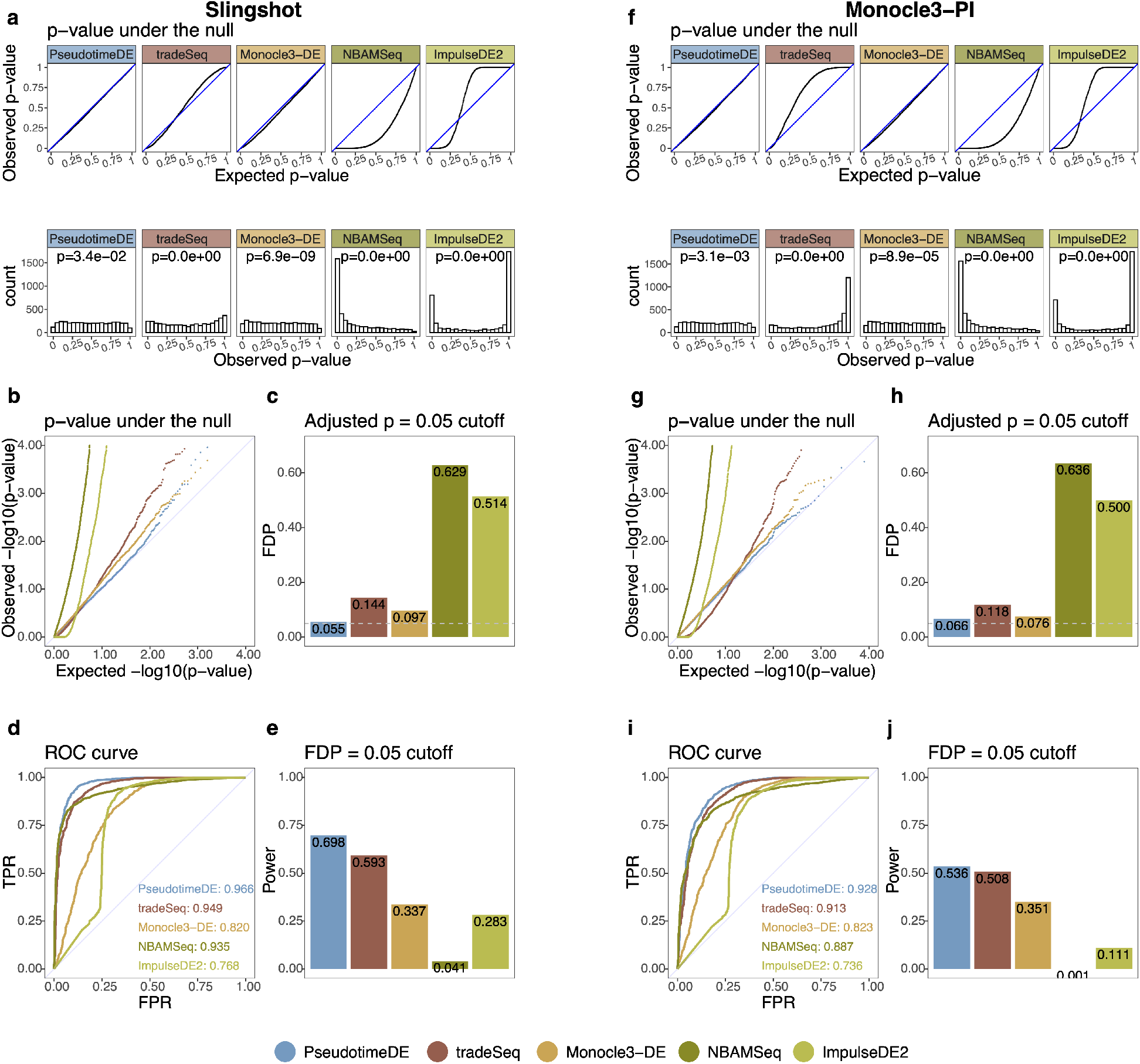
PseudotimeDE outperforms four state-of-the-art methods (tradeSeq, Monocle3-DE, NBAMSeq, and ImpulseDE2) for identifying DE genes along cell pseudotime. Left panels (a)–(e) are based on pseudotime inferred by Slingshot; right panels (f)–(j) are based on pseudotime inferred by Monocle3-PI. **(a) & (f)** Distributions of non-DE genes’ observed *p*-values by five DE methods with inferred pseudotime. Top: quantile-quantile plots that compare the empirical quantiles of the observed *p*-values against the expected quantiles of the Uniform[0,1] distribution. Bottom: histograms of the observed *p*-values. The *p*-values shown on top of histograms are from the Kolmogorov–Smirnov test under the null hypothesis that the distribution is Uniform[0,1]. The larger the *p*-value, the more uniform the distribution is. Among the five DE methods, PseudotimeDE’s observed *p*-values follow most closely the expected Uniform[0,1] distribution. **(b) & (g)** Quantile-quantile plots of the same *p*-values as in (a) and (f) on the negative log_10_ scale. PseudotimeDE returns better-calibrated small *p*-values than the other four methods do. **(c) & (h)** FDPs of the five DE methods with the target FDR 0.05 (BH adjusted-*p* ≤0.05). PseudotimeDE yields the FDP closest to 0.05. **(d) & (i)** ROC curves and AUROC values of the five DE methods. PseudotimeDE achieves the highest AUROC. **(e) & (j)** Power of the five DE methods under the FDP = 0.05 cutoff. PseudotimeDE achieves the highest power.

Next, we compare these methods in terms of their ability to distinguish DE genes from non-DE genes, ability measured by the area under the receiver operating characteristic curve (AUROC) values (Fig. 3d & i and Additional file 1: Figs. S3–S5d & i). PseudotimeDE achieves the highest AUROC values. Among the other four methods, tradeSeq and NBAMSeq have slightly lower AUROC values than PseudotimeDE’s, and Monocle3-DE and ImpulseDE2 have much lower AUROC values than the other three methods’. The reason is that PseudotimeDE, tradeSeq, and NBAMSeq all use the flexible model NB-GAM, while Monocle3-DE and ImpulseDE2 use much more restrictive models, which limit their power.

Realizing that the ill-calibrated *p*-values of the existing four methods invalidate their FDR control, we compare all five methods in terms of their power under an actual 5% false discovery proportion (FDP, defined as the proportion of false discoveries among the discoveries in one synthetic dataset) instead of the nominal 5% FDR. Our results show that PseudotimeDE achieves the highest power on all datasets except for the bifurcation dataset, where PseudotimeDE has slightly lower power than tradeSeq’s (Fig. 3e & j and Additional file 1: Figs. S3–S5e & j). These results demonstrate the high power of PseudotimeDE and its effective FDR control, which is lacking in existing methods.

In summary, our simulation results verify that PseudotimeDE outperforms existing methods in terms of generating well-calibrated *p*-values, which are essential for FDR control, and identifying DE genes with high power. Notably, the two bulk RNA-seq methods, NBAMSeq and ImpulseDE2, yield worse results than the three scRNA-seq methods do. Hence, we only focus on the scRNA-seq methods in the following three real data applications.

### Real data example 1: dendritic cells stimulated with lipopolysaccharide

In the first application, we compare PseudotimeDE with tradeSeq and Monocle3-DE on a dataset of mouse dendritic cells (DCs) after stimulation with lipopolysaccharide (LPS, a component of gram-negative bacteria) [31]. In this dataset, gene expression changes are expected to be associated with the immune response process. We first apply Slingshot and Monocle3-PI to this dataset to infer cell pseudotime, and then we input the inferred pseudotime into PseudotimeDE, tradeSeq, and Monocle3-DE for DE gene identification. Consistent with our simulation results, the *p*-values of tradeSeq are ill-calibrated: their bimodal distributions indicate that they do not follow the uniform distribution under the null hypothesis; instead, many of them are inflated, and this inflation would lead to power loss in DE gene identification (Fig. 4a & e). Indeed, at a nominal Benjamini-Hochberg (BH) adjusted *p*-value ≤ 0.01 threshold (which corresponds to controlling the FDR ≤ 1% when *p*-values are valid), tradeSeq identifies the smallest number of DE genes, while PseudotimeDE identifies the most DE genes, followed by Monocle3-DE. Notably, most of the DE genes identified by tradeSeq are also identified by PseudotimeDE (Fig. 4b & f), a result consistent with the over-conservativeness of tradeSeq due to its inflated *p*-values. Unlike tradeSeq, Monocle3-DE does not exhibit the inflated *p*-value issue; however, it uses a more restrictive model than PseudotimeDE and tradeSeq do. Hence, we use functional analyses to investigate whether Monocle3-DE misses certain DE genes due to its restrictive modeling. We also investigate whether the additional DE genes found by PseudotimeDE but missed by tradeSeq or Monocle3-DE are biologically meaningful.

**Figure 4:**
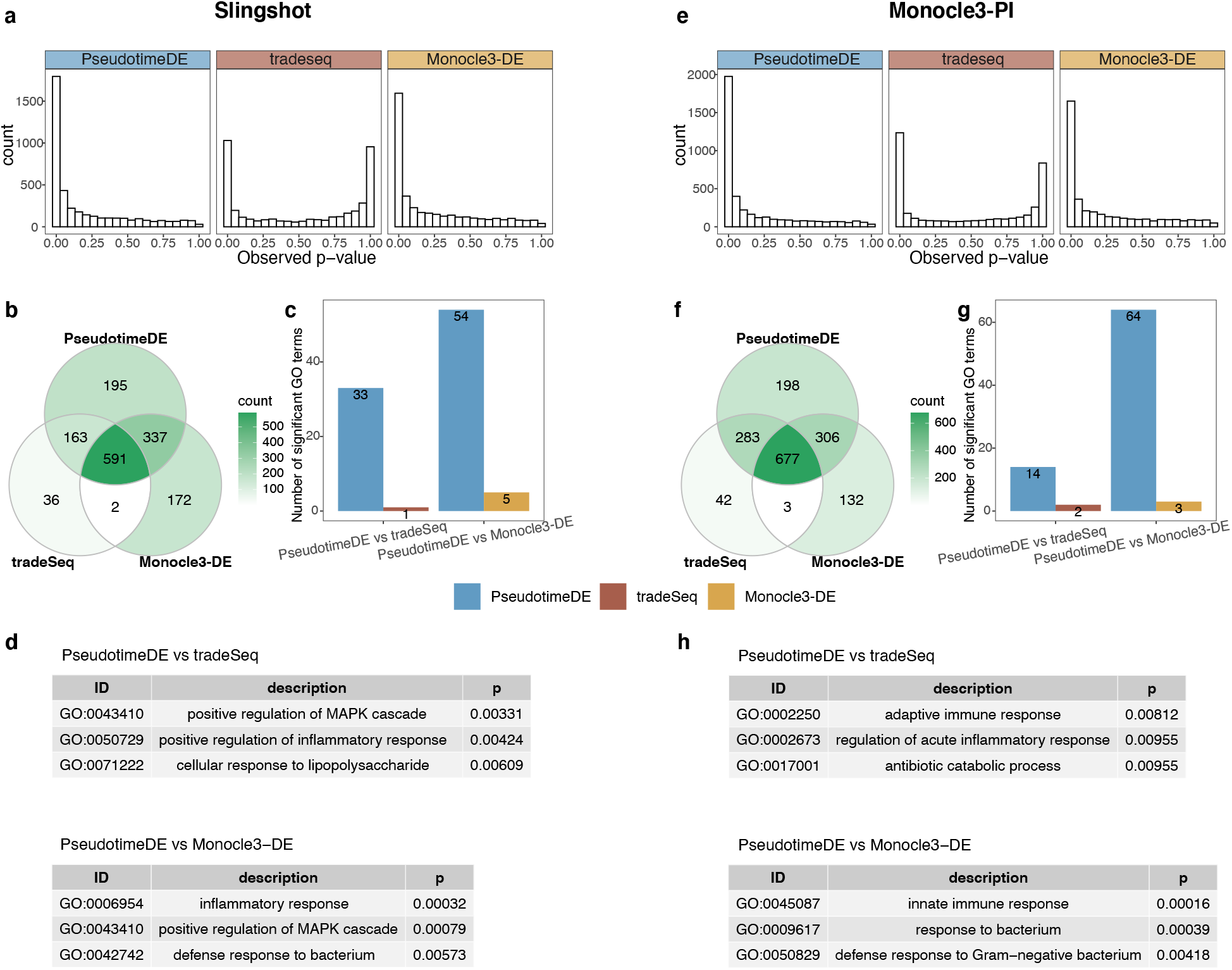
Application of PseudotimeDE, tradeSeq, and Monocle3-DE to the LPS-dendritic cell dataset. Left panels (a)–(d) are based on pseudotime inferred by Slingshot; right panels (e)–(h) are based on pseudotime inferred by Monocle3-PI. **(a) & (e)** Histograms of all genes’ *p*-values by the three DE methods. The bimodal distributions of tradeSeq’s *p*-values suggest a violation of the requirement that *p*-values follow the Uniform[0,1] distribution under the null hypothesis. **(b) & (f)** Venn plots showing the overlaps of the significant DE genes (BH adjusted-*p* ≤ 0.01) identified by the three DE methods. PseudotimeDE’s DE genes nearly include tradeSeq’s. **(c) & (g)** Numbers of GO terms enriched (*p* < 0.01) in the significant DE genes specifically found by PseudotimeDE or tradeSeq/Monocle3-DE in pairwise comparisons between PseudotimeDE and tradeSeq/Monocle3-DE in (b) & (f). Many more GO terms are enriched in the PseudotimeDE-specific DE genes than in the tradeSeq- or Monocle3-DE-specific ones. **(d) & (h)** Example GO terms enriched in the Pseudotime-specific DE genes in (c) & (g). Many of these terms are related to LPS, immune process, and defense to bacterium.

Our first strategy is to perform gene ontology (GO) analysis on the DE genes identified by each method and compare the enriched GO terms. We find that more GO terms are enriched (with enrichment *p*-values < 0.01) in the DE genes identified by PseudotimeDE (Additional file 1: Fig. S6a & c), and that the PseudotimeDE-specific GO terms are related to immune responses (Additional file 1: Fig. S6b & d). However, comparing enriched GO terms does not directly reflect the difference of DE genes identified by different methods. Hence, our second strategy is to probe the functions of the DE genes that are uniquely identified by one method in pairwise comparisons of PseudotimeDE vs. tradeSeq and PseudotimeDE vs. Monocle3-DE. We first perform GO analysis on each set of uniquely identified DE genes. For a fair comparison of two methods, we remove the overlapping DE genes found by both methods from the background gene list in GO analysis. Our results show that many more GO terms are enriched (with enrichment *p*-values < 0.01) in Pseudotime-specific DE genes than in tradeSeq- or Monocle3-DE-specific DE genes (Fig. 4c & g). Moreover, many of those PseudotimeDE-specific GO terms are directly related to the immune responses of DCs to LPS stimulation, including the GO terms “cellular response to lipopolysaccharide” and “defense response to Gram-negative bacterium” (Fig. 4d & h; Additional file 2: Table S1). To focus more on immune responses, we next perform enrichment analysis using the immunologic signatures (C7) in the Molecular Signatures Database (MSigDB) [32]. Our results show that only PseudotimeDE-specific DE genes have enriched MSigDB C7 terms (with BH adjusted p values < 0.01), while tradeSeq- and Monocle3-DE-specific DE genes have almost no enrichment (Additional file 1: Fig. S7a & c). More importantly, many enriched terms in PseudotimeDE-specific DE genes were found by previous studies of DCs stimulated with LPS (see examples in Additional file 1: Fig. S7b & d; Additional file 2: Table S1); this is direct evidence that supports the validity of PseudotimeDE-specific DE genes. For illustration purpose, we visualize the expression levels of some known and novel DE genes identified by PseudotimeDE using UMAP, and clear DE patterns are observed (Additional file 1: Fig. S8–S9). In conclusion, our functional analyses verify that PseudotimeDE identifies biologically meaningful DE genes missed by tradeSeq and Monocle3-DE, confirming that PseudotimeDE has high power in addition to its well-calibrated *p*-values.

### Real data example 2: pancreatic beta cell maturation

In the second application, we compare PseudotimeDE with tradeSeq and Monocle3-DE on a dataset of mouse beta cell maturation process [33]. We first apply Slingshot and Monocle3-PI to this dataset to infer cell pseudotime, and then we input the inferred pseudotime into PseudotimeDE, tradeSeq, and Monocle3-DE for DE gene identification. Consistent with previous results, the *p*-values of tradeSeq follow a bimodal distribution, suggesting that many of them are incorrectly inflated (Fig. 5a & f). At the nominal BH-adjusted *p*-value ≤ 0.01 level, PseudotimeDE identifies the second most DE genes, fewer than Monocle3-DE’s identified DE genes and much more than tradeSeq’s (Fig. 5b & g). As the numbers of identified DE genes cannot reflect these methods’ performance, we use three approaches to evaluate the DE genes identified by each method.

**Figure 5:**
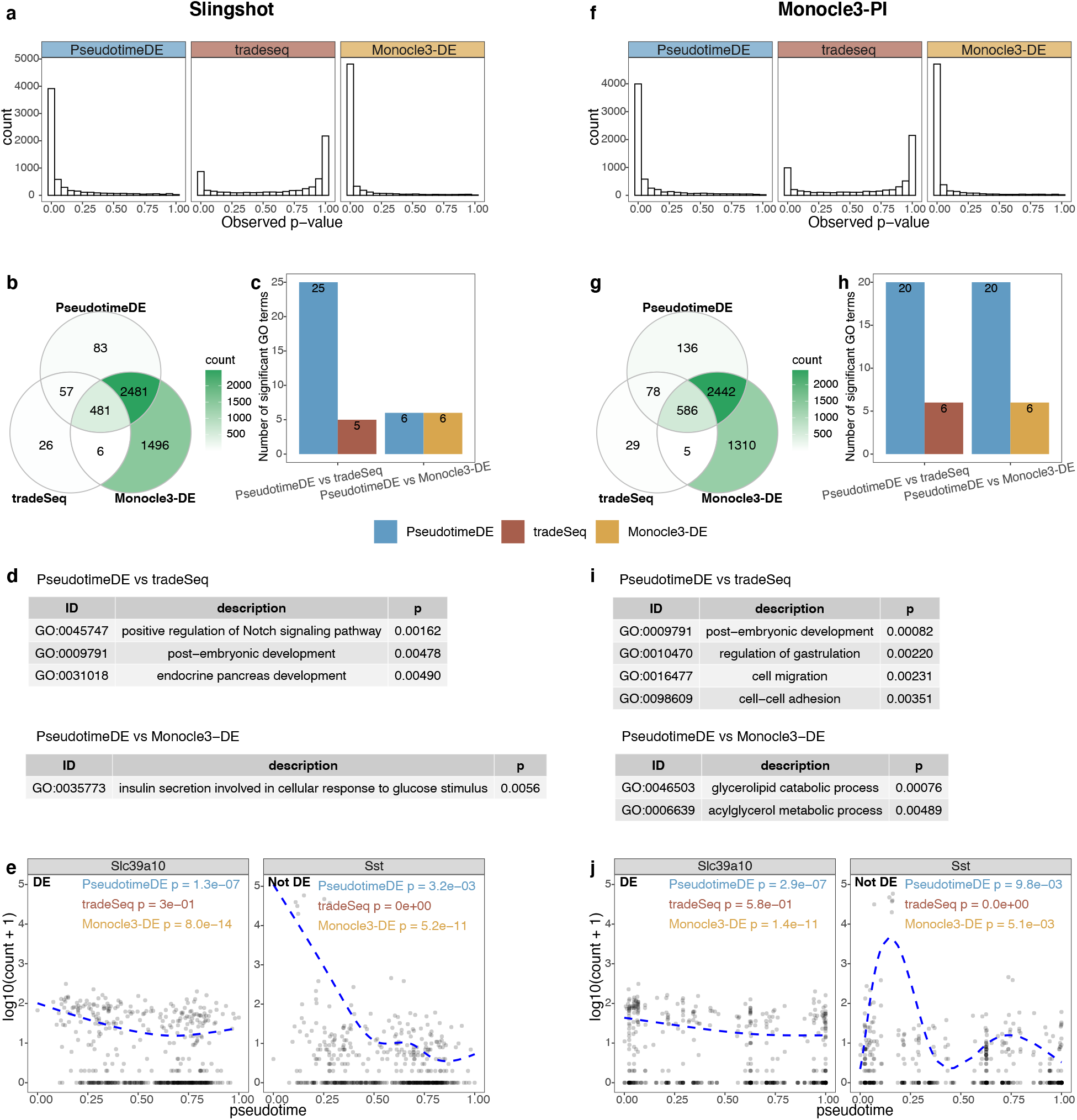
Application of PseudotimeDE, tradeSeq, and Monocle3-DE to the pancreatic beta cell maturation dataset. Left panels (a)–(e) are based on pseudotime inferred by Slingshot; right panels (f)–(j) are based on pseudotime inferred by Monocle3-PI. **(a) & (f)** Histograms of all genes’ *p*-values by the three DE methods. The bimodal distributions of tradeSeq’s *p*-values suggest a violation of the requirement that *p*-values follow the Uniform[0,1] distribution under the null hypothesis. **(b) & (g)** Venn plots showing the overlaps of the significant DE genes (BH adjusted-*p* ≤ 0.01) identified by the three DE methods. PseudotimeDE’s DE genes nearly include tradeSeq’s. **(c) & (h)** Numbers of GO terms enriched (*p* < 0.01) in the significant DE genes specifically found by PseudotimeDE or tradeSeq/Monocle3-DE in pairwise comparisons between PseudotimeDE and tradeSeq/Monocle3-DE in (b) & (g). Many more GO terms are enriched in the PseudotimeDE-specific DE genes than in the tradeSeq- or Monocle3-DE-specific ones. **(d) & (i)** Example GO terms enriched in the Pseudotime-specific DE genes in (c) & (h). Many of these terms are related to related to insulin, beta cell regulation, and pancreas development. **(e) & (j)** Two examples genes: *Slc39a10* (DE) and *Sst* (non-DE). For *Slc39a10*, both PseudotimeDE and Monocle3-DE yield small *p*-values (*p* < 1*e* − 6), while tradeSeq does not (*p* > 0.1). For Sst, PseudotimeDE yields larger *p*-values than tradeSeq and Monocle3-DE do. Dashed blue lines are the fitted curves by NB-GAM.

We first perform GO analysis on each set of uniquely identified DE genes, using the same pairwise comparisons of PseudotimeDE vs. tradeSeq and PseudotimeDE vs. Monocle3-DE as for the LPS-dendritic data. Our results show that more GO terms are enriched (with enrichment *p*-values < 0.01) in PseudotimeDE-specific DE genes than in tradeSeq- or Monocle3-DE-specific DE genes (Fig. 5c & h). Moreover, many of those PseudotimeDE-specific GO terms are directly related to pancreatic beta cell development, e.g., “positive/negative regulation of Notch signaling pathway”[34] and “endocrine pancreas development” (Fig. 5c & h; Additional file 3: Table S2). As a complementary result, we also perform GO analysis on the DE genes identified by each method. We find that the GO terms, which are only enriched in the DE genes identified by PseudotimeDE, are related to beta cell development and thus more biologically meaningful than the GO terms that are only enriched in the DE genes identified by tradeSeq or Monocle3-DE (Additional file 1: Fig. S10b & d; Additional file 3: Table S2).

Second, we utilize the DE genes identified from bulk RNA-seq data in the original paper [33] to evaluate the DE gene rankings established by PseudotimeDE, tradeSeq, and Monocle3-DE from scRNA-seq data. Taking the bulk DE genes as a gene set, we perform the geneset enrichment analysis (GSEA) [32] on all genes’ – log_10_ *p*-values output by PseudotimeDE, tradeSeq, and Monocle3-DE. Among the three methods, PseudotimeDE leads to the highest normalized enrichment score (NES) (Additional file 3: Table S2), suggesting that the bulk DE genes are most enriched in the top-ranked DE genes found by PseudotimeDE.

Third, we examine a highly credible DE gene *Slc39a10* [33, 35] and a verified non-DE gene *Sst* [33] as representative examples. For *Slc39a10*, both PseudotimeDE and Monocle3-DE yield small *p*-values (< 10^−6^), while tradeSeq outputs a *p*-value > 0.1 and thus misses it (Fig. 5e & g). For Sst, PseudotimeDE yields the largest *p*-value (> 0.001), while tradeSeq and Monocle3-DE yield extremely small *p*-values (< 10^−10^) and thus mistaken it as a DE gene. Hence, PseudotimeDE has the best performance on these two representative genes.

For illustration purpose, we visualize the expression levels of some known and novel DE genes identified by PseudotimeDE using UMAP, and clear DE patterns are observed (Additional file 1: Figs. S11–S12).

### Real data example 3: bone marrow differentiation

In the third application, we compare PseudotimeDE with tradeSeq and Monocle3-DE on a dataset of mouse bone marrow differentiation [36]. We apply Slingshot with UMAP for dimensionality reduction to infer cell pseudotime as described in the tradeSeq paper [16]. Slingshot constructs the reported bifurcation topology (in the tradeSeq paper) on the original dataset (Fig. 6a), but it infers trifurcation topology, instead of bifurcation topology, on 40% of subsamples (Fig. 6b shows randomly picked ten subsamples). Note that the third lineage consisting of the cell type megakaryocyte (MK) was reported in the Monocle2 paper ([12]), suggesting the observed topology uncertainty may be biologically meaningful.

**Figure 6:**
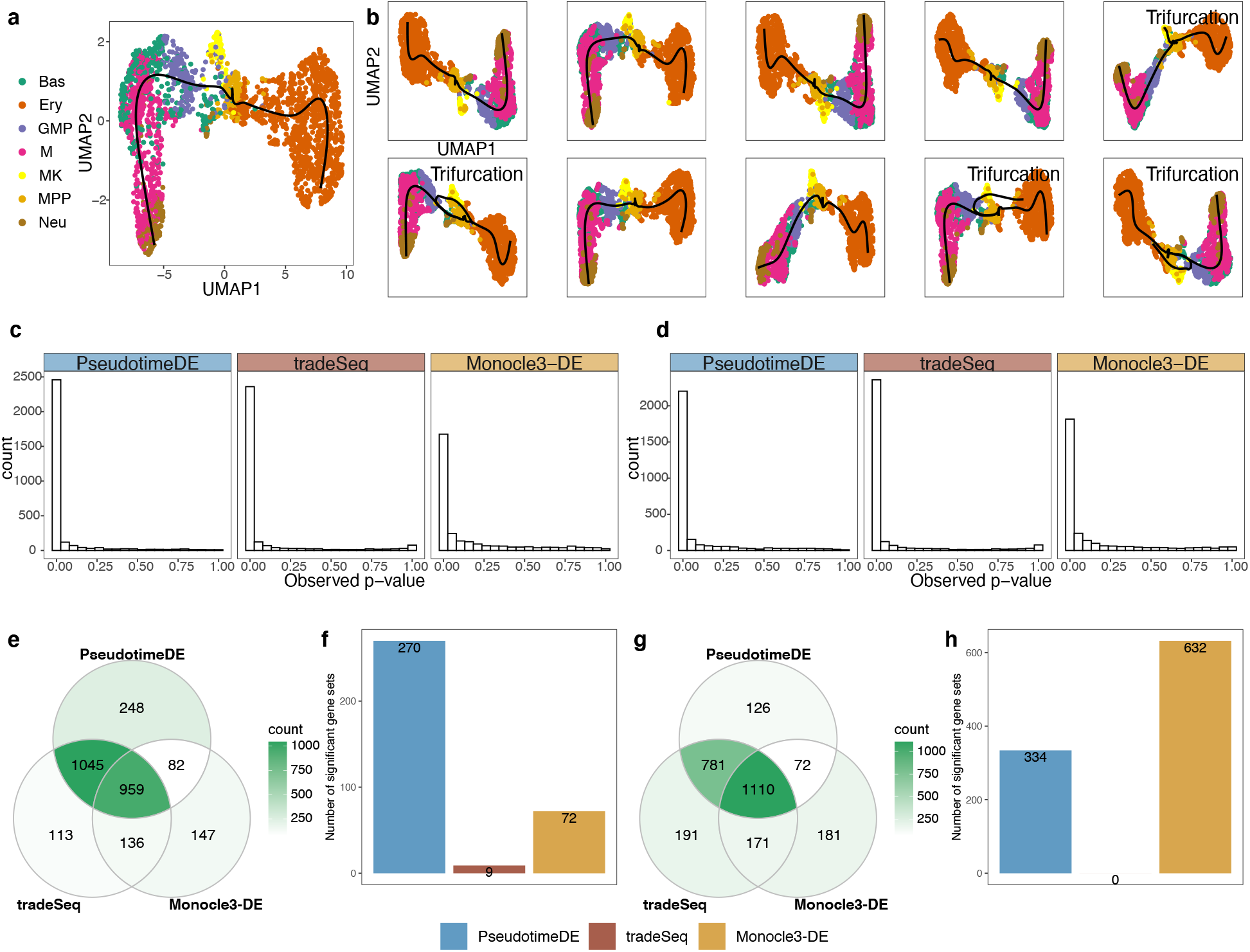
Application of PseudotimeDE, tradeSeq, and Monocle3-DE to the mouse bone marrow dataset. **(a)** UMAP visualization and inferred pseudotime by Slingshot. Pre-defined cell types are marked by colors. Slingshot returns a bifurcation topology, denoted as lineage 1 (left) and lineage 2 (right). **(b)** UMAP visualization and inferred pseudotime by Slingshot on ten random subsamples. Four out of ten subsamples do not yield bifurcation topology but trifurcation topology, where the third lineage mainly contains the cell type “MK” and was reported in [12]. **(c)** Histograms of all genes’ *p*-values calculated by the three DE methods in the first lineage. **(d)** Histograms of all genes’ *p*-values calculated by the three DE methods in the second lineage. **(e)** Venn plot showing the overlaps of the significant DE genes (BH adjusted-*p* ≤ 0.01) identified by the three DE methods in lineage 1. PseudotimeDE and tradeSeq share 77.6% (Jaccard index) DE genes. **(f)** Numbers of enriched gene sets (*q* < 0.25) by GSEA using the *p*-values in lineage 1 by the three DE methods. Although the DE genes are similar in (e), PseudotimeDE yields 270 enriched gene sets, while tradeSeq only yields 9. **(g)** Venn plot showing the overlaps of the significant DE genes (BH adjusted-*p* ≤ 0.01) identified by the three DE methods in lineage 2. Similar to lineage 1 in (g), PseudotimeDE and tradeSeq share 77.2% (Jaccard index) DE genes. **(h)** Numbers of enriched gene sets (*q* < 0.25) by GSEA using the *p*-values in lineage 2 by the three DE methods. PseudotimeDE and Monocle3-DE yield hundreds of enriched gene sets, while tradeSeq does not yield any enriched gene sets.

For a fair comparison, we only make pseudotimeDE use the subsamples with inferred bifurcation topology, because both tradeSeq and Monocle3-DE use the inferred bifurcation topology from the original data to identify DE genes. Consistent with previous results, the tradeSeq *p*-values follow a bimodal distribution that is unexpected for well-calibrated *p*-values. At a nominal BH-adjusted *p*-value ≤ 0.01 threshold, the three methods identify highly similar DE genes (Fig. 6e & g). For instance, PseudotimeDE and tradeSeq share about 80% of their identified DE genes (Jaccard index). From the few method-specific DE genes, functional analyses cannot indicate which method performs better. Therefore, we use GSEA instead to evaluate methods’ *p*-values. Surprisingly, although the three methods identify highly similar DE genes, their *p*-values lead to vastly different GSEA results. At the q < 0.25 level, PseudotimeDE and Monocle3-DE yield hundreds of enriched gene sets, while tradeSeq only yields a few or no enriched gene sets (Fig. 6f & h; Additional file 4: Table S3). This result indicates that, besides the ranking of *p*-values, the nominal values of *p*-values are also crucial for downstream analysis such as GSEA. Hence, the well-calibrated *p*-values make PseudotimeDE superior to existing methods for DE gene identification and downstream analyses.

### Real data example 4: natural killer T cell subtypes

In the fourth application, we compare PseudotimeDE with tradeSeq and Monocle3-DE on a dataset of natural killer T cell (NKT cell) subtypes [37]. We apply Slingshot with PCA for dimensionality reduction to infer cell pseudotime and construct the trifurcation topology (Fig. 7a) reported in the original study. We apply the three DE methods to identify DE genes in each of the three lineages. Consistent with the previous results, the *p*-values of tradeSeq follow a bimodal distribution, suggesting that many of them are incorrectly inflated (Fig. 7b).

**Figure 7:**
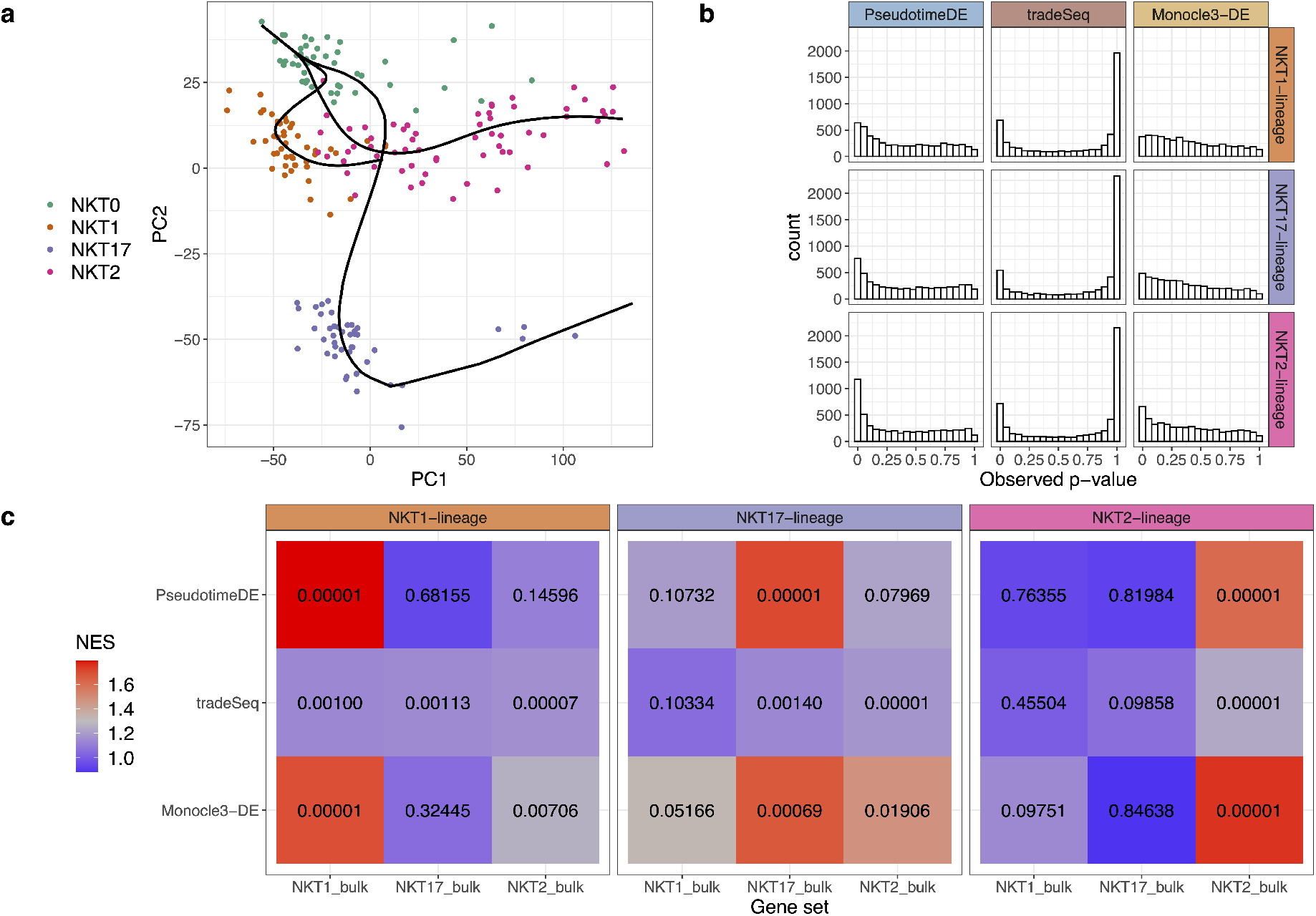
Application of PseudotimeDE, tradeSeq, and Monocle3-DE to the natural killer T cell dataset. **(a)** PCA visualization and inferred pseudotime by Slingshot. Pre-defined NKT subtypes are marked by colors. Slingshot returns a trifurcation topology, where the three lineages are NKT0 to NKT1, NKT0 to NKT17, and NKT0 to NKT2. **(b)** Histograms of all genes’ *p*-values in the three lineages calculated by the three DE methods. **(c)** Heatmaps of normalized enrichment scores (NESs, marked by colors) and their corresponding *p*-values (in numbers) from the GSEA. Each NES value and its corresponding *p*-value are calculated for each DE method and each lineage, based on the *p*-values of a DE method for a lineage and that lineage’s DE genes found from bulk RNA-seq data, denoted by “NKT1 bulk”, “NKT17 bulk,” or “NKT2 bulk” [37]. Note that among the three DE methods, PseudotimeDE outputs *p*-values that best agree with the lineage-specific DE genes from bulk data and thus most distinguish the three lineages. For instance, for the NKT1 lineage, PseudotimeDE’s small *p*-values are enriched in the “NKT1 bulk” gene set only, while tradeSeq and Monocle3-DE have small *p*-values enriched in at least two lineage-specific DE gene sets.

For validation purpose, we utilize the lineage-specific DE genes identified from bulk RNA-seq data in the original study [37] to evaluate the DE gene rankings established by PseudotimeDE, tradeSeq, and Monocle3-DE from scRNA-seq data. Specifically, we perform the GSEA using the bulk DE gene sets in the same way as for the pancreatic beta cell maturation dataset. The GSEA shows that PseudotimeDE’s *p*-values best agree with the lineage-specific DE genes from bulk data and thus most distinguish the three lineages. For example, for the NKT1 lineage, PseudotimeDE’s small *p*-values are exclusively enriched in the “NKT1 bulk” gene set, while tradeSeq and Monocle3-DE have small *p*-values enriched in at least two lineage-specific DE gene sets (Fig. 7c). This result confirms that, compared with the DE genes identified by the other two DE methods, the top DE genes identified by PseudotimeDE are more biologically meaningful.

### Real data example 5: cell cycle phases

In the fifth application, we compare PseudotimeDE with tradeSeq and Monocle3-DE on a dataset of human induced pluripotent stem cells (iPSCs) measured with cell cycle phases (FUCCI labels) [38]. The original study has reported 101 cyclic genes whose expression levels have large proportions of variance explained (PVE) explained by cells’ FUCCI labels [38]; that is, cells’ FUCCI labels are regarded as the predictor, a gene’s expression levels in the same cells are regarded as the response, and a PVE is calculated from a nonparametric smoothing fit; hence, the larger the PVE, the better the gene’s expression levels can be predicted by the cell cycle phases. The original study has also developed an R package `peco` to infer cell cycle phases from scRNA-seq data.

In our study, we first construct a benchmark dataset by treating the 101 cyclic genes as true DE genes and using the same genes with expression levels randomly shuffled across cells as the true non-DE genes; hence, our positive and negative sets both contain 101 genes. Then we apply the R package `peco` to this dataset to infer each cell’s cycle phase, which is equivalent to pseudotime; that is, we use `peco` as the pseudotime inference method. Finally, we apply the three DE methods.

Our results show that, for the true non-DE genes, only PseudotimeDE generates valid *p*-values that approximately follow the Uniform[0,1] distribution (Fig. 8a & c). For the true DE genes, PseudotimeDE’s (– log_10_ transformed) *p*-values, one per gene, have the highest correlation with these genes’ PVE, indicating that PseudotimeDE successfully identifies the top DE genes as those with the strongest cyclic trends (Fig. 8b). PseudotimeDE also yields successful FDR control, the highest AUROC value, and the highest power, among the three DE methods (Fig. 8d, e and f). Therefore, we conclude that PseudotimeDE outperforms tradeSeq and Monocle3-DE in identifying cell cycle-related genes from this iPSC scRNA-seq dataset.

**Figure 8:**
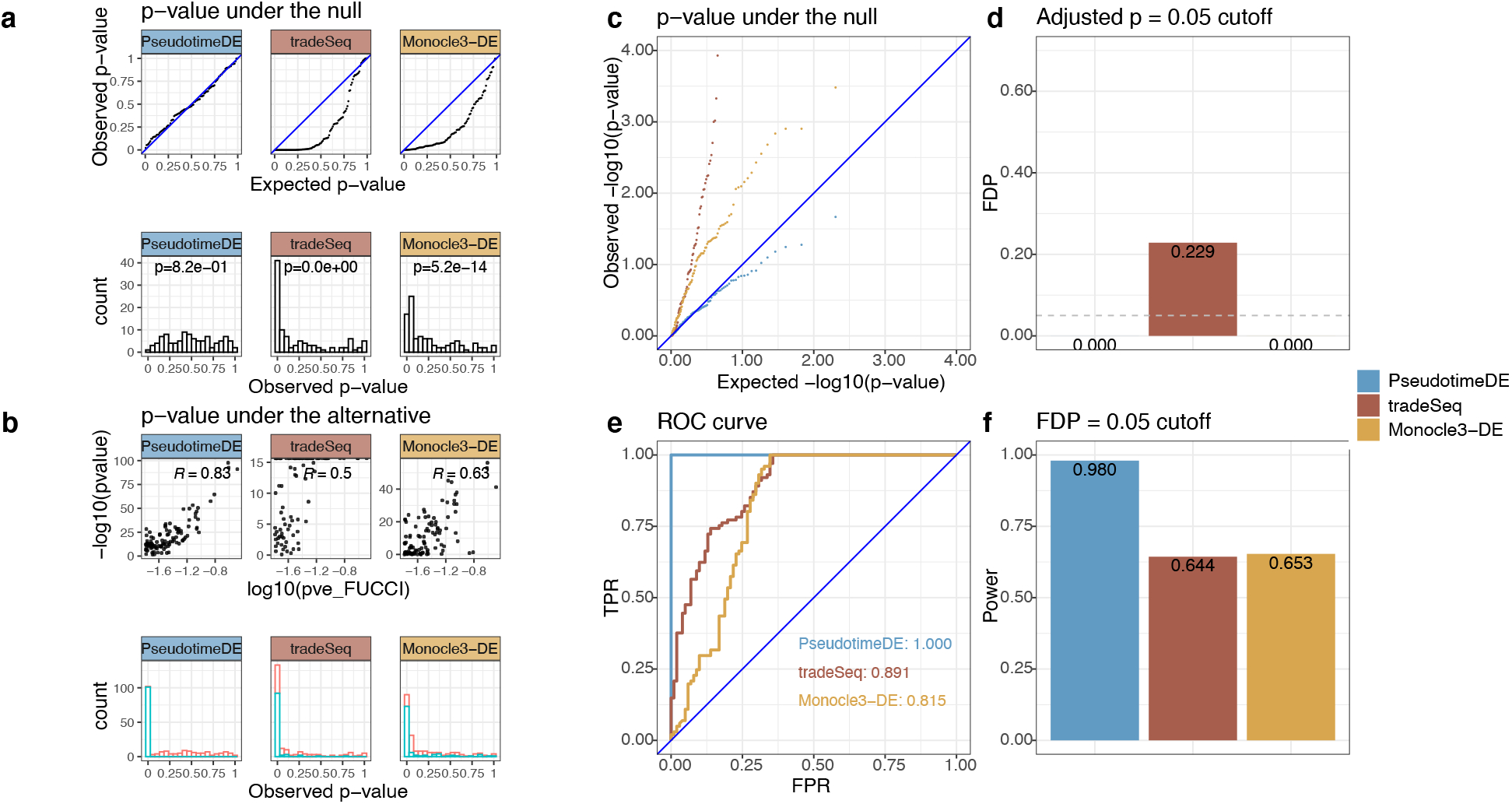
Application of PseudotimeDE, tradeSeq, and Monocle3-DE to the cell cycle phase dataset. **(a)** Distributions of non-DE genes’ *p*-values by three DE methods with inferred pseudotime. Top: quantile-quantile plots that compare the empirical quantiles of non-DE genes’ *p*-values against the expected quantiles of the Uniform[0,1] distribution. Bottom: histograms of non-DE genes’ *p*-values. The *p*-values shown on top of histograms are from the Kolmogorov–Smirnov test under the null hypothesis that the distribution is Uniform[0,1]. The larger the *p*-value, the more uniform the distribution is. Among the three DE methods, PseudotimeDE’s *p*-values follow most closely the expected Uniform[0,1] distribution. **(b)** Distributions of DE genes’ *p*-values by three DE methods with inferred pseudotime. Top: scatter plots of DE genes’ *p*-values against the proportions of variance explained (PVE), which measure the strengths of genes’ inferred cyclic trends in the original study [38]. PseudotimeDE’s *p*-values (– log_10_ transformed) have the highest correlation with the PVE, indicating that PseudotimeDE identifies the genes with the strongest cyclic trends as the top DE genes. Bottom: histograms of all genes’ *p*-values. Blue and red colors represent the *p*-values of DE genes and non-DE genes (same as in (a) bottom), respectively. PseudotimeDE yields the best separation of the two gene groups’ *p*-values. **(c)** Quantile-quantile plots of the same *p*-values as in (a) on the negative log_10_ scale. PseudotimeDE returns the best-calibrated *p*-values. **(d)** FDPs of the three DE methods with the target FDR 0.05 (BH adjusted-*p* ≤ 0.05). **(e)** ROC curves and AUROC values of the three DE methods. PseudotimeDE achieves the highest AUROC. **(f)** Power of the three DE methods under the FDP = 0.05 cutoff. PseudotimeDE achieves the highest power.

### PseudotimeDE allows users to inspect the uncertainty of inferred cell pseudotime

Besides the identification of DE genes, PseudotimeDE offers functionality for inspecting the uncertainty of pseudotime inference via its intermediate subsampling step. In Figs. 2e & f and 6a & b, PseudotimeDE reveals the uncertainty of inferred cell lineages. Users usually want to fix the lineage topology, e.g., a bifurcation topology, for downstream analysis; however, the topology can vary across subsamples. Hence, we recommend that users check if the inferred topology from the original data can also be inferred from more than half of the subsamples. If not, users may consider using another pseudotime inference method with less uncertainty or adding additional constraints on inferred topology. Otherwise, the great uncertainty of inferred cell lineage would impair the reliability of downstream analyses.

Next, conditioning on a given lineage topology, PseudotimeDE allows users to visualize the uncertainty of pseudotime within a lineage (Fig. 2a–d), also guiding the choice of pseudotime inference methods in terms of uncertainty.

### Computational time

The only feasible way to accommodate all pseudotime inference methods and to account for their uncertainty in DE gene identification is to use subsampling and permutation, the approach taken by PseudotimeDE. However, a common concern of permutation-based methods is that they are computationally intensive. Admittedly, PseudotimeDE is slower than existing non-permutation-based methods, but its computational time is nevertheless acceptable to server users. With 24 cores (Intel ”Cascade Lake” CPU), 36 GB RAM and 1000 subsamples, PseudotimeDE takes 3–8 hours to analyze each of the first three scRNA-seq datasets in our study. Specifically, the LPS-dendritic cell dataset (4016 genes, 390 cells) takes 3 hours, the pancreatic beta cell maturation dataset (6121 genes, 497 cells) takes 3.5 hours, and the bone marrow dataset (3004 genes, 2660 cells, two lineages) takes 8 hours. The computational time is proportional to the number of genes, the number of lineages, and the number of subsamples. Of course, it is inversely proportional to the number of available cores.

To reduce the computational time of PseudotimeDE, users have two options. First, they may reduce the number of genes to be tested. For instance, lowly expressed genes, such as those with more than 90% of zero counts, are recommended to be filtered out because they are usually of less interest to biologists. Second, they may reduce the number of subsamples. Due to its parametric estimation of the null distribution of the test statistic, PseudotimeDE does not require an enormous number of subsamples. We find that PseudotimeDE with only 100 subsamples generates similar *p*-values to those based on 1000 subsamples (Additional file 1: Fig. S20). If using 100 subsmaples, the computational time is within 0.5 hour.

In an undesirable scenario that computational resources are too limited, users have to abandon the consideration of pseudotime uncertainty and treat inferred pseudotime as fixed. Then they do not need the subsampling procedure, and PseudotimeDE will calculate *p*-values from the asymptotic null distribution of the test statistic [39], with a short computational time as other nonpermutation-based methods’.

## Discussion

We propose a statistical method PseudotimeDE to identify DE genes along inferred cell pseudotime. PseudotimeDE focuses on generating well-calibrated *p*-values while taking into account the randomness of inferred pseudotime. To achieve these goals, PseudotimeDE first uses subsampling to estimate the uncertainty of pseudotime. Second, PseudotimeDE fits the NB-GAM or ZINB-GAM to both the original dataset and the permuted subsampled datasets to calculate the test statistic and its approximate null values. Next, PseudotimeDE fits a parametric distribution to estimate the approximate null distribution of the test statistic. Finally, PseudotimeDE calculates *p*-values with a high resolution. PseudotimeDE is flexible to accommodate cell pseudotime inferred in a standard format by any method. Its use of NB-GAM and ZINB-GAM allows it to capture diverse gene expression dynamics and to accommodate undesirable zero inflation in data.

Comprehensive studies on simulated and real data confirm that PseudotimeDE yields better FDR control and higher power than four existing methods (tradeSeq, Monocle3-DE, NBAMSeq, and ImpulseDE2) do. On simulation data, PseudotimeDE generates well-calibrated *p*-values that follow the uniform distribution under the null hypothesis, while existing methods except Monocle3-DE have *p*-values violating the uniform assumption. Well-calibrated *p*-values guarantee the valid FDR control of PseudotimeDE. Moreover, thanks to its use of flexible models NB-GAM and ZINB-GAM, PseudotimeDE has higher power than Monocle3-DE, which uses a more restrictive model GLM and thus has less power. PseudotimeDE also outperforms the other three methods–tradeSeq, NBAMSeq, and ImpulseDE2—that generate ill-calibrated *p*-values in terms of power. On three real scRNA-seq datasets, the DE genes uniquely identified by PseudotimeDE embrace better biological interpretability revealed by functional analyses, and the *p*-values of PseudotimeDE lead to more significant GSEA results.

An interesting and open question is what pseudotime inference method works the best with PseudotimeDE. While we observe that PseudotimeDE has higher power with Slingshot than with Monocle3-PI in simulation studies, we realize that the reason may be associated with the simulation design (e.g., the lineage structures), and thus we cannot draw a conclusion from this observation. Due to the diversity of biological systems and the complexity of pseudotime inference [9], we decide to leave the choice of pseudotime inference methods open to users, and this is the advantage of PseudotimeDE being flexible to accommodate inferred pseudotime by any methods. In practice, we encourage users to try popular pseudotime inference methods and use PseudotimeDE as a downstream step to identify DE genes, so that they can analyze the identification results and decide which pseudotime inference method is more appropriate for their dataset.

The zero inflation, or “dropout” issue, remains perplexing and controversial in the single-cell field [40–44]. The controversy is regarding whether excess zeros that cannot be explained by Poisson or negative binomial distributions are biological meaningful or not. Facing this controversy, we provide two models in PseudotimeDE: NB-GAM and ZINB-GAM, the former treating excess zeros as biologically meaningful and the latter not. Specifically, the negative binomial distribution in NB-GAM is fitted to all gene expression counts including excess zeros, while the fitting of the negative distribution in ZINB-GAM excludes excess zeros, which ZINB-GAM implicitly treats as non-biological zeros. PseudotimeDE allows the choice between the two models to be user specified or data driven. From our data analysis, we realize that the choice often requires biological knowledge of the dataset to be analyzed. Specifically, on the LPS-dendritic cell dataset and pancreatic beta cell maturation dataset, we observe that ZINB-GAM leads to power loss: some potential DE genes cannot be identified by ZINB-GAM because zero counts contain useful information (Additional file 1: Figs. S15–S18). Our observation is consistent with another recent study [42], whose authors observed that “zero-inflated models lead to higher false-negative rates than identical non-zero-inflated models.” Hence, our real data analysis results are based on NB-GAM. However, realizing the complexity of biological systems and scRNA-seq protocols, we leave the choice between NB-GAM and ZINB-GAM as an option for users of PseudotimeDE, and we encourage users to plot their known DE genes as in Additional file 1: Figs. S15–S18 to decide which of NB-GAM and ZINB-GAM better captures the gene expression dynamics of their interest.

The current implementation of PseudotimeDE is restricted to identifying the DE genes that have expression changes within a cell lineage. While methods including GPfates [45], Monocle2 BEAM [46], and tradeSeq can test whether a gene’s expression change is associated with a branching event leading to two lineages, they do not consider the uncertainty of lineage inference. How to account for such topology uncertainty is a challenging open question, as we have seen in Figs. 2f and 6b that the inferred lineage may vary from a bifurcation topology to a trifurcation topology on different subsets of cells. A possible direction is to use the selective inference [47, 48], and we will leave the investigation of this question to future research. Due to this topology uncertainty issue, PseudotimeDE is most suitable for single-cell gene expression data that contain only one cell lineage (including cyclic data) or a small number of well separated cell lineages (e.g., bifurcation). The reason is that these data can maintain stable inferred cell topology after subsampling, an essential requirement of PseudotimeDE. That said, PseudotimeDE is not designed for data with many equivocal cell lineages or a complex cell hierarchy, the data that cannot maintain stable inferred cell topology across subsamples, because for such data, it is difficult to find one-to-one matches between cell lineages inferred from a subsample and those inferred from the original data. Then, a practical solution for such data is to first define a cell lineage of interest and then apply PseudotimeDE to only the cells assigned to this lineage.

There are other open questions to be explored. An important question is: when do we want to identify DE genes along pseudotime? As we have shown in the **Results** section, inferred pseudotime can be highly uncertain. As biologists often sequence cells at multiple physical time points if they want to investigate a biological process, a straightforward analysis is to identify the DE genes that have expression changes across the physical time points. Then we have two definitions of DE genes: the genes whose expression changes across pseudotime vs. physical time. Understanding which definition is more biologically relevant is an open question. Another question is whether it is possible to integrate pseudotime with physical time to identify biologically relevant DE genes. Answering either question requires a statistical formulation that is directly connected to a biological question.

Another question is how to explore gene-gene correlations along cell pseudotime. Current DE methods only detect marginal gene expression changes but ignore gene-gene correlations. It remains unclear whether gene-gene correlations are stable or varying along cell pseudotime. Hence, a new statistical method to detect gene-gene correlation changes along inferred pseudotime may offer new biological insights into gene co-expression and regulation at the single-cell resolution.

## Methods

### PseudotimeDE methodology

Here we describe the methodological detail of PseudotimeDE, a pseudotime-based differential expression testing method. As an overview, PseudotimeDE takes a scRNA-seq count matrix and a pseudotime inference method as input, estimates the uncertainty of pseudotime, performs a differential expression test, and returns a *p*-value of each gene. The core of PseudotimeDE is to obtain a valid null distribution of the DE gene test statistic so that the *p*-values are well-calibrated.

### Mathematical notations

We denote by **Y** = (*Y_ij_*) an *n* × *m* gene expression count matrix, whose rows and columns correspond to *n* cells and *m* genes, respectively; that is, *Y_ij_* is the read count of gene *j* in cell *i*. Taking **Y** as input, a pseudotime inference method would return a pseudotime vector **T** = (*T*_1_,…, *T_i_*,…, *T_n_*)^T^, where *T_i_*, ∈ [0,1] denotes the normalized inferred pseudotime of cell *i* (i.e., the cells with the smallest and largest pseudotime have *T_i_* = 0 and 1, respectively; normalization is used for visualization simplicity). Note that *T_i_*, is a random variable due to the random-sampling nature of the *n* cells and the possible uncertainty introduced by the pseudotime inference method.

### Uncertainty estimation

To estimate the uncertainty of pseudotime **T**, we subsample 80% cells (rows) in **Y** for *B* times. Although there are some theoretical results about the optimal subsample size [49], they do not apply to our problem setting. Hence, we simply choose 80% because it is widely used [50, 51], similar to the popularity of 5-fold cross validation in machine learning [52]. Simulation results also supports that 80% is a reasonable choice, and PseudotimeDE is robust to various subsampling proportions (Additional file 1: Fig. S22; see Additional file 1 for detail). It is worth noting that the bootstrap technique is inapplicable for our problem because it leads to repeated sampling of the same cell, causing issues with some pseudotime inference methods such as Monocle2. If the cells have pre-defined groups (i.e., cell types), we use the stratified sampling by first subsampling 80% cells within each group and then combining these within-group subsamples into one subsample. By default, we set *B* = 1000. For each subsample 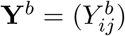, an *n′* × *m* matrix where *n′* = ⌊.8*n*⌋, we perform pseudotime inference with the same parameters used for the original dataset **Y**. As a result, we obtain *B* subsample-based realizations of pseudotime **T**: {**T**^1^, ···, **T**^*b*^, ···, **T**^*B*^}, where **T**^*b*^ ∈ [0,1]^*n′*^, and each cell appears in approximately 80% of these *B* realizations. Note that we have to apply pseudotime inference to each subsample before permutation to account for pseudotime inference uncertainty; otherwise, if each subsample’s pseudotime is just a subsample of all cells’ pseudotime, we are essentially treating all cells’pseudotime as fixed, and the uncertainty in pseudotime inference is ignored. Here is the mathematical explanation. Given that we have *n* cells with inferred pseudotime as *T*_1_,…, *T_n_*, if we use direct subsampling, then in the *b*-th subsampling, the subsampled *n′* cells’ pseudotime is just a size-*n′* subsample of {*T*_1_,…, *T*_*n*_}. Instead, in PseudotimeDE, the subsampled *n′* cells’ inferred pseudotime 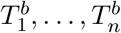, may be *n′* values that are not in *T*_1_,…, *T_n_*. In other words, the uncertainty in pseudotime inference is reflected in 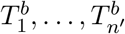.

### PseudotimeDE model

We use the negative binomial–generalized additive model (NB-GAM) as the baseline model to describe the relationship between every gene’s expression in a cell and the cell’s pseudotime. For gene *j* (*j* = 1,…, *m*), its expression *Y_ij_* in cell *i* and the pseudotime *T_i_* of cell *i* (*i* = 1,…, *n*) are assumed to follow

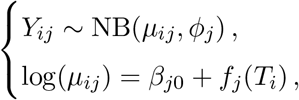

where NB(*μ_ij_*, *ϕ_j_*) denotes the negative binomial distribution with mean *μ_ij_* and dispersion *ϕ_j_*, and 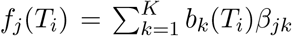 is a cubic spline function. The number of knots *k* is predefined as 6 and usually has little effect on results [53]. For gene *j*, PseudotimeDE fits the NB-GAM to (*Y*_1*j*_,…, *Y_nj_*)^T^ and **T** = (*T*_1_,…, *T_n_*)^T^ using the R package `mgcv` (version 1.8.31), which estimates model parameters by penalized-restricted maximum likelihood estimation.

To account for excess zeros in scRNA-seq data that may not be explained by the NB-GAM, we introduce a hidden variable *Z_ij_* to indicate the “dropout” event of gene *j* in cell *i*, and the resulting model is called the zero-inflated negative binomial–generalized additive model (ZINB-GAM):

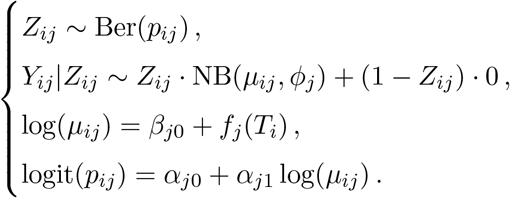

For gene *j*, PseudotimeDE fits the ZINB-GAM to (*Y*_1*j*_,…, *Y_nj_*)^T^ and **T** = (*T*_1_,…, *T_n_*)^T^ using the expectation-maximization (EM) algorithm, which is partially based on R package `zigam` [54]. To use PseudotimeDE, users can specify whether to use the ZINB-GAM or NB-GAM. If users do not provide a specification, PseudotimeDE will automatically choose between the two models for each gene by the Akaike information criterion (AIC). By default, PseudotimeDE uses NB-GAM unless the AIC of ZINB-GAM exceeds the AIC of NB-GAM by at least 10, a threshold suggested by [55].

### Statistical test and *p*-value calculation

To test if gene *j* is DE along cell pseudotime, PseudotimeDE defines the null and alternative hypotheses as

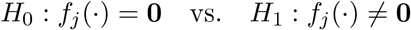

We denote the estimate of (*f_j_*(*T*_1_),…, *f_j_*(*T_n_*))^T^ by 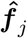, whose estimated covariance matrix (of dimensions *n* × *n*) is denoted by 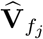. Then the test statistic is

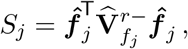

where 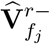 is the rank-*r* pseudoinverse of 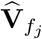, where *r* is determined in the way described in [39]. When the *T_i_*’s are fixed, the asymptotic null distribution of *S_j_* is described in [39], and the *p*-value can be calculated by the R package `mgcv`.

A key novelty of PseudotimeDE is its accounting for the uncertainty of inferred pseudotime. When the *T_i_*’s are random, the asymptotic null distribution of *S_j_* given that *T_i_*’s are fixed [39] and the *p*-value calculation in the R package `mgcv` no longer apply. To address this issue and estimate the null distribution, PseudotimeDE uses the following permutation procedure: (1) PseudotimeDE randomly permutes each subsample-based realization 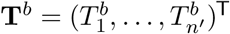 into 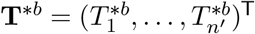 (2) PseudotimeDE fits the above model to 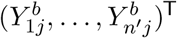 and **T**^\**b*^, and calculates the test statistic *S_j_*’s value as 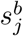 using the R package `mgcv`; (3) PseudotimeDE performs (1) and (2) for *b* = 1,…, *B* and collects the resulting 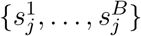 as the null values of the test statistic *S_j_*.

Then PseudotimeDE estimates the null distribution of *S_j_* in two ways. Based on the estimated null distribution in either way and the observed test statistic value *s_j_*, which is calculated from the original dataset by the R package `mgcv`, PseudotimeDE calculates a *p*-value for gene *j*.

1. **Empirical estimate**. PseudotimeDE uses the empirical distribution of 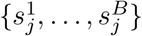 as the estimated null distribution. Following the suggestion in [56], PseudotimeDE calculates the *p*-value of gene *j* as

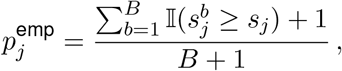

where 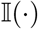 is the indicator function. We refer to this *p*-value as the “empirical *p*-value.”
2. **Parametric estimate**. The resolution of 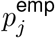 depends on the number of permutations *B*, because the smallest value 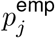 may take is 1/(*B* + 1). Although users often cannot afford a too large *B* due to limited computational resources, they still desire a high resolution of *p*-values to control the FDR to a small value (e.g., 5%) when the number of tests (i.e., the number of genes in DE gene identification) is large. To increase the resolution of *p*-values, PseudotimeDE fits a parametric distribution to 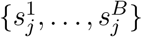 and uses the fitted distribution as the estimated null distribution. Driven by the empirical distribution of 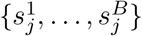, PseudotimeDE considers two parametric distributions: (1) a gamma distribution Γ(*α, β*) with *α, β* > 0 and (2) a two-component gamma mixture model *γ*Γ(*α*_1_, *β*_1_) + (1 − *γ*)Γ(*α*_2_, *β*_2_) with 0 < *γ* < 1 and *α*_1_, *β*_1_, *α*_2_, *β*_2_ > 0. After fitting both distributions to 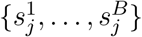 using the maximum likelihood estimation (gamma distribution fit by the R package `fitdistrpius` (version 1.0.14) [57] and gamma mixture model fit by the R package `mixtools` (version 5.4.5) [58]), PseudotimeDE chooses between the two fitted distributions by performing the likelihood ratio test (LRT) with 3 degrees of freedom (i.e., difference in the numbers of parameters between the two distributions). If the LRT *p*-value is less or equal than 0.01, PseudotimeDE uses the fitted two-component gamma mixture model as the parametric estimate of the null distribution of *S_j_*; otherwise, PseudotimeDE uses the fitted gamma distribution. The Anderson-Darling goodness-of-fit test verifies that such a parametric approach fits the empirical distributions well (Additional file 1: Fig. S21; see Additional file 1 for detail). Denoting the cumulative distribution function of the parametrically estimated null distribution by 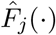, PseudotimeDE calculates the *p*-value of gene *j* as

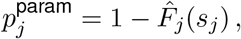

where is referred to as the “parametric *p*-value.” PseudotimeDE outputs both 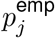 and 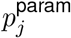 for gene *j, j* = 1,…, *m*. Empirical evidence shows that parametric *p*-values agree with empirical *p*-values well across the [0, 1] interval (Additional file 1: Fig. S19). All the findings in the Results section are based on 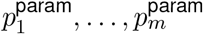 due to their higher resolution.

### Pseudotime inference methods

We apply two state-of-the-art methods, Slingshot and Monocle3-PI, to inferring the cell pseudotime of each dataset. For single-lineage data, we specify the start cluster in Slingshot and the start node in Monocle3-PI. For bifurcation/trifurcation data, we specify the start cluster/node and the end clusters/nodes in Slingshot/Monocle3-PI. By default, the dimensionality reduction methods used for pseudotime inference are PCA and UMAP for Slingshot and Monocle3-PI, respectively. The R Bioconductor package `slingshot` (version 1.4.0) and the R package `monocle3` (version 0.2.0) are used.

### DE analysis methods

We compare PseudotimeDE with four existing methods for identifying DE genes along pseudotime/time-course from scRNA-seq data (tradeSeq and Monocle3-DE) or bulk RNA-seq data (ImpulseDE2 and NBAMSeq). All these methods take a count matrix **Y** and a pseudotime vector **T** as input, and they return a *p*-value for each gene. For tradeSeq, we use the functions `fitGAM` and `associationTest` (https://statomics.github.io/tradeSeq/articles/tradeSeq.html). The number of knots parameter *K* in tradeSeq is chosen by 100 random genes based on the tradeSeq vignette. For Monocle3-DE, we use the function `fit_models` (https://cole-trapnell-lab.github.io/monocle3/docs/differential/). Since ImpulseDE2 cannot be applied to scRNA-seq data directly, we follow the modified implementation of ImpulseDE2 in the tradeSeq paper (https://github.com/statOmics/tradeSeqPaper). The R Bioconductor packages `tradeSeq` (version 1.3.15), `monocle3` (version 0.2.0), `ImpulseDE2` (version 1.10.0), and `NBAMSeq` (version 1.10.0) are used.

### Functional (gene ontology and gene-set enrichment) analyses

We use the R package `topGO` (version 2.38.1) [59] to perform the gene-ontology (GO) enrichment analysis on identified DE genes. We use the R package `clusterprofiler` (version 3.14.3) [60] to perform the gene-set enrichment analysis (GSEA) analysis on ranked gene lists, where genes in each list are ranked by their ranking sores defined as – log_10_ transformed *p*-values (the gene with the smallest *p*-value is ranked the top); *p*-values that are exactly zeros are replaced by one-tenth of the smallest non-zero *p*-value. If unspecified, the GO terms are “biological process (BP)” terms.

### Simulation study

We use the R package `dyntoy` (0.9.9) to generate single-lineage data and bifurcation data. For single-lineage data, we generate three datasets with increasing dispersion levels (low dispersion, medium dispersion, and high dispersion). Each single-lineage dataset consists of 500 cells and 5000 genes (with 20% as DE genes). For bifurcation data, we use the medium dispersion level. The bifurcation dataset consists of 750 cells and 5000 genes (with 20% as DE genes).

### Case studies

#### LPS-dendritic cell dataset

this Smart-seq dataset contains primary mouse dendritic cells (DCs) stimulated with lipopolysaccharide (LPS) [31], available at Gene Expression Omnibus (GEO) under accession ID GSE45719. In our analysis, we use the the cells from 1h, 2h, 4h, and 6h in the pre-processed data from the study that benchmarked pseudotime inference methods [9]. After the genes with > 90% zeros are removed, the final dataset consists of 4016 genes and 390 cells, which are expected to be in a single lineage. When applying tradeSeq, we use the recommended ZINB-WaVE [61] + tradeSeq procedure to account for potential zero-inflation. The R Bioconductor package `zinbwave` (version 1.8.0) is used.

#### Pancreatic beta cell maturation dataset

this Smart-seq2 dataset measures the maturation process of mouse pancreatic beta cells [33], available at GEO under accession ID GSE87375. We use the cells from cell type “beta” in the pre-processed data from the study that benchmarked pseudotime inference methods [9]. After the genes with > 90% zeros are removed, the final dataset consists of 6121 genes and 497 cells, which are expected to be in a single lineage. When applying tradeSeq, we use the recommended ZINB-WaVE + tradeSeq procedure to account for potential zero-inflation. The R Bioconductor package `zinbwave` (version 1.8.0) is used.

#### Mouse bone marrow dataset

this MARS-seq dataset contains myeloid progenitors in mouse bone marrow [36], available at GEO under accession ID GSE72859. We use the pre-processed data provided by the tradeSeq vignette. After the genes with > 90% zeros are removed, the final dataset consists of 3004 genes and 2660 cells. We follow the procedure of combining UMAP and Slingshot to infer pseudotime as described in tradeSeq paper [16]

#### Natural killer T cell dataset

this Smart-seq2 dataset measures four natural killer T cell (NKT cell) subtypes in mouse [37], available at GEO under accession ID GSE74597. We use the preprocessed data from the study that benchmarked pseudotime inference methods [9]. After the genes with > 90% zeros are removed, the final dataset consists of 5270 genes and 197 cells, which are expected to have three lineages. We use PCA + Slingshot to infer the pseudotime. When applying tradeSeq, we use the recommended ZINB-WaVE + tradeSeq procedure to account for potential zero-inflation. The R Bioconductor package `zinbwave` (version 1.8.0) is used.

#### Cell cycle phase dataset

this Fluidigm protocol dataset measures human induced pluripotent stem cells (iPSCs) [38]. The iPSCs are FUCCI-expressing so that their cell cycle phases can be tracked. The authors also developed an R package `peco` for predicting cell cycle phases from single-cell gene expression data. We use the example dataset provided by `peco`, which consists of 101 known cell cycle-related genes (DE genes). To construct null cases, we randomly shuffle the 101 DE genes’ expression levels across cells to create 101 non-DE genes. The final dataset consists of 202 genes and 888 cells. We use the R package `peco` (version 1.1.21) to infer cell cycle phases.

## Availability of data and materials

The R package `PseudotimeDE` is available at https://github.com/SONGDONGYUAN1994/PseudotimeDE[62] under the MIT license. The source code and data for reproducing the results are available at http://doi.org/10.5281/zenodo.4663580 [63].

## Acknowledgements

The authors would like to thank Dr. Kelly Street for his suggestions on the usage of Slingshot and uncertainty estimate. The authors would like to thank Mr. Huy Nguyen for his assistance in simulation. The authors also appreciate the comments and feedback from the members of the Junction of Statistics and Biology at UCLA (http://jsb.ucla.edu).

## Funding

This work was supported by the following grants: National Science Foundation DBI-1846216, NIH/NIGMS R01GM120507, Johnson & Johnson WiSTEM2D Award, Sloan Research Fellowship, and UCLA David Geffen School of Medicine W.M. Keck Foundation Junior Faculty Award (to J.J.L.).

## Author information

### Authors Contributions

D.S. and J.J.L. designed the research. D.S. conducted the research and wrote the computer code. J.J.L. supervised the project. D.S. and J.J.L. discussed the results and wrote the manuscript.

### Corresponding author

Correspondence to Jingyi Jessica Li.

## Ethics declarations

### Ethics approval and consent to participate

None.

### Consent for publication

Not applicable for this study.

### Competing interests

None.

## Additional files

### Additional file 1

Supplementary materials. It includes all supplementary text and figures.

### Additional file 2

Supplementary table 1. It includes enrichment analysis results of LPS-dendritic cell dataset.

### Additional file 3

Supplementary table 2. It includes enrichment analysis results of pancreatic beta cell maturation dataset.

### Additional file 4

Supplementary table 3. It includes enrichment analysis results of mouse bone marrow dataset.

## Supplementary Materials

### Goodness-of-fit of the parametric distribution

We perform a goodness-of-fit test to show that our parametric approach fits the empirical distributions well (Fig. S21). We choose the Anderson-Darling (AD) test since it gives more weights on the tail of the distribution than the Kolmogorov-Smirnov test does. On one simulated dataset and one real dataset, the distribution of AD test *p*-values is approximately Uniform[0,1](Fig. S21a & c), indicating that the parametric distributions fit the empirical null distributions well (i.e., all cases are from the null). Since this goodness-of-fit check is crucial, `PseudotimeDE` R package automatically outputs the AD test *p*-value per gene so that users can check whether the parametric approximation works well; if not, users may increase the number of subsampling and use the empirical null for DE *p*-value calculation.

### Choice of the subsampling proportion

In general, results of PseudotimeDE is robust to the subsampling proportion. We have applied to one simulation dataset (Fig. S22) five sampling proportions (50%, 60%, 70%, 80%, and 90%), among which 80% is the default used in PseudotimeDE. From the results, we first observe that the *p*-values (on the – log_10_ scale) are highly correlated between 80% and any other proportion (Fig. S22c), confirming the robustness of PseudotimeDE to the subsampling proportion. Second, we observe that the power of PseudotimeDE remains stable under varying sampling proportions, though the power is slightly lower when the sampling proportion is 50% (Fig. S22a). Third, the false discovery proportion (FDP) of PseudotimeDE is also stable under varying sampling proportions, though the FDP is slightly higher than the nominal level when the sampling proportion is 90% (Fig. S22b). Therefore, we conclude that 80% is a reasonable default sampling proportion.

### Differences of the NB-GAM fitting in PseudotimeDE, tradeSeq, and NBAMSeq

PseudotimeDE, tradeSeq, and NBAMSeq all rely on the R package `mgcv` for fitting NB-GAM by penalized-restricted maximum likelihood. Therefore, the model fitting of NB-GAM is overall similar in the three methods. There are only minor differences, listed below.

1. In PseudotimeDE, the dispersion parameter *ϕ_j_* of gene *j* is estimated simultaneously with the mean parameter *μ_ij_* of gene *j* in cell *i* by the penalized-restricted maximum likelihood (PRML); in contrast, NBAMSeq uses the maximum a posterior (MAP) dispersion estimate by R package `DESeq2` [21]. NBAMSeq is designed for bulk RNA-seq and accounts for a samll sample size (i.e., number of replicates); therefore, the MAP estimate by `DESeq2` is more biased (for variance reduction purpose) than the PRML estimate used by PseudotimeDE. Given that scRNA-seq has a large enough sample size (i.e., number of cells), we think that the PRML estimate of *ϕ_j_* is preferred over the MAP estimate.
2. In PseudotimeDE, the number of knots *K* is pre-defined as a fix number (the default *K* = 6, which is usually large enough); In contrast, tradeSeq randomly subsamples a small set of genes to choose a *K* for all genes.
3. PseudotimeDE fits a NB-GAM to each lineage, while tradeSeq incorporates a lineage covariate so that it can fit a NB-GAM to multiple lineages jointly. The reason is that PseudotimeDE needs to account for the uncertainty in pseudotime inference and within-lineage model fitting makes the inference easier.
4. Pseudotime also implements a zero-inflated NB-GAM (ZINB-GAM) that includes a dropout probability parameter for each gene, and PseudotimeDE estimates this parameter internally. In contrast, while tradeSeq can also turn its NB-GAM into ZINB-GAM, it estimates the dropout probability parameters externally by ZINB-WaVE [61]; NBAMSeq is designed for bulk RNA-seq data and does not consider zero inflation.

Although the NB-GAM fitting of PseudotimeDE is similar to that of the other two methods, its inference (p-value calculation) is totally different, and this is the main novelty of PseudotimeDE.

### The number of knots *K*

The number of knots, *K*, a parameter in the generalized additive model (GAM), defines the number and positions of piecewise polynomial functions in a spline function.

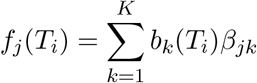

where *f_j_*(·) is the smooth spline function of gene *j*, *T_i_* is the pseudotime of cell *i*, and *K* is the number of knots. It is worth noting that *K* − 1 is also the **nominal degree of freedom** of the model.

*K* needs to be specified before the negative binomial-GAM (NB-GAM) is fitted. For the GAM model, “exact choice of *K* is not generally critical [64],” if the GAM model is correctly used for inference. This is why we do not include the *K* selection step in PseudotimeDE. The default *K* = 6 in PseudotimeDE is usually large enough for modelling gene trajectories. Using a larger *K* is not harmful, but it will increase the computational time.

In contrast, tradeSeq has a *K* selection step, which we think is problematic. To choose *K*, tradeSeq randomly samples a small set of genes and uses AIC to find an “appropriate”K for all genes (see *Choosing an appropriate number of knots* in the tradeSeq paper [16]). We follow their vignette and obtain *K* = 4 in Fig. 3a and *K* = 6 in Fig. 3f. In tradeSeq, the null distribution of the test statistic *S_j_* is

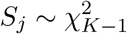

Therefore, when *K* = 4 (Fig. 3a), the null distribution of every gene is 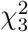; when *K* = 6 (Fig. 3f), the null distribution of every gene is 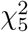. As a result, the distributions of tradeSeq *p*-values under the null are quite different between Fig. 3a and Fig. 3f. It seems like the results of tradeSeq is highly sensitive to the choice of *K*. Such sensitivity is probably due to their incorrect null distribution. That is, tradeSeq uses the nominal degree of freedom *K* − 1 in their null distribution 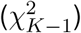; however, the correct way is to use the effective degree of freedom, which is gene-specific and always smaller than *K* − 1 due to the penalization on the likelihood function. As a result, tradeSeq uses null distributions (of its test statistics) that have heavier right tails than those of the correct null distributions, resulting in conservative *p*-values (whose distribution under the null has a mode near 1). Details about the theory and usage of the effective degree of freedom in GAM can be found in Dr. Simon Woods’ book [53] and 2013 *Biometrika* paper [39].

**Figure S1:**
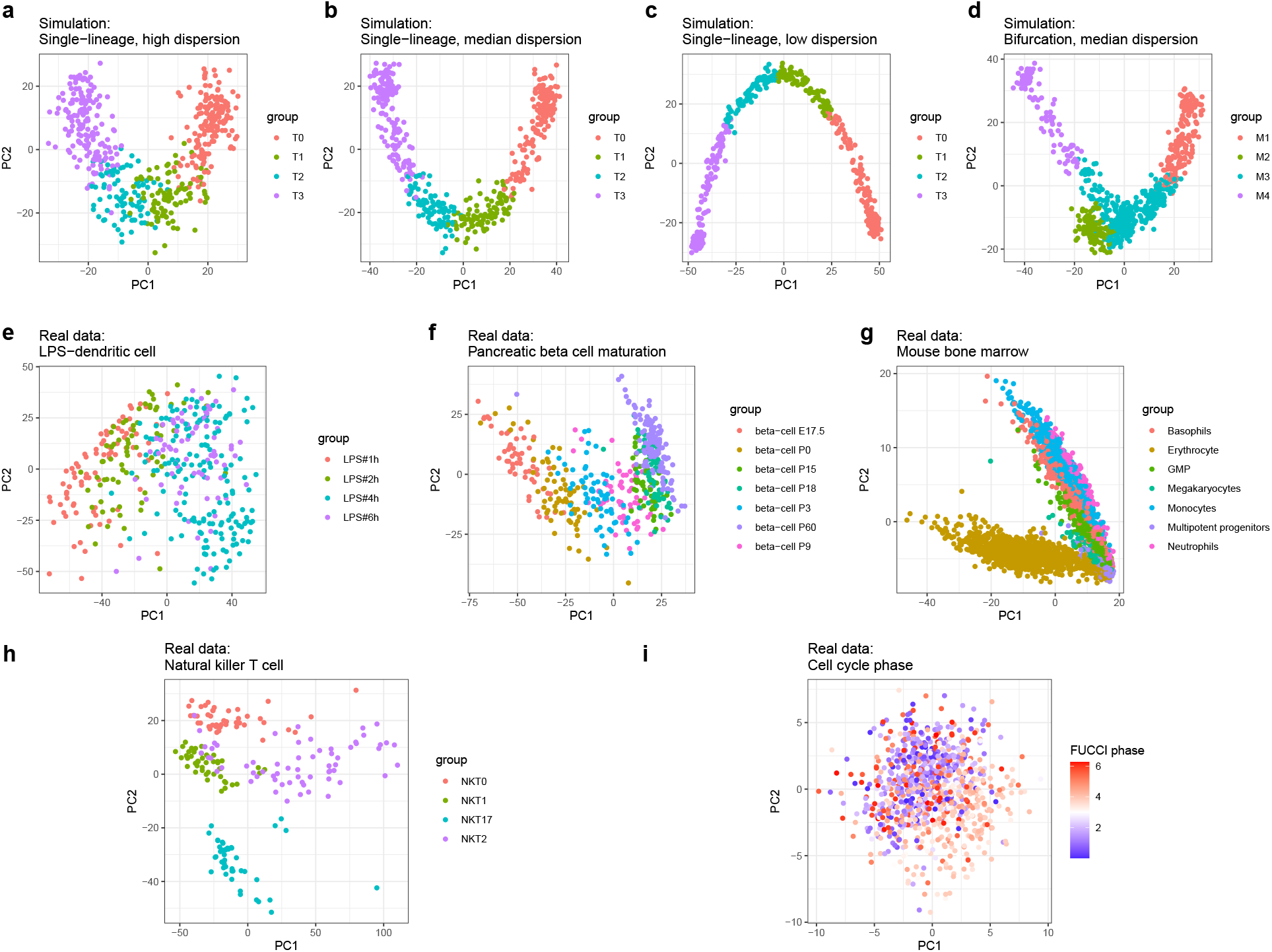
PCA visualization of datasets. PCA visualization of all synthetic and real datasets used in this paper. Panels (a)–(d) are synthetic datasets, where the groups correspond to time points. Panels (e)–(h) are real datasets, where the groups correspond to time points in (e) & (f) or annotated cell types in (g) & (h). Panel (i) is a real dataset with external cell cycle information, where the FUCCI phase indicates experimentally measured cell cycle phase.

**Figure S2:**
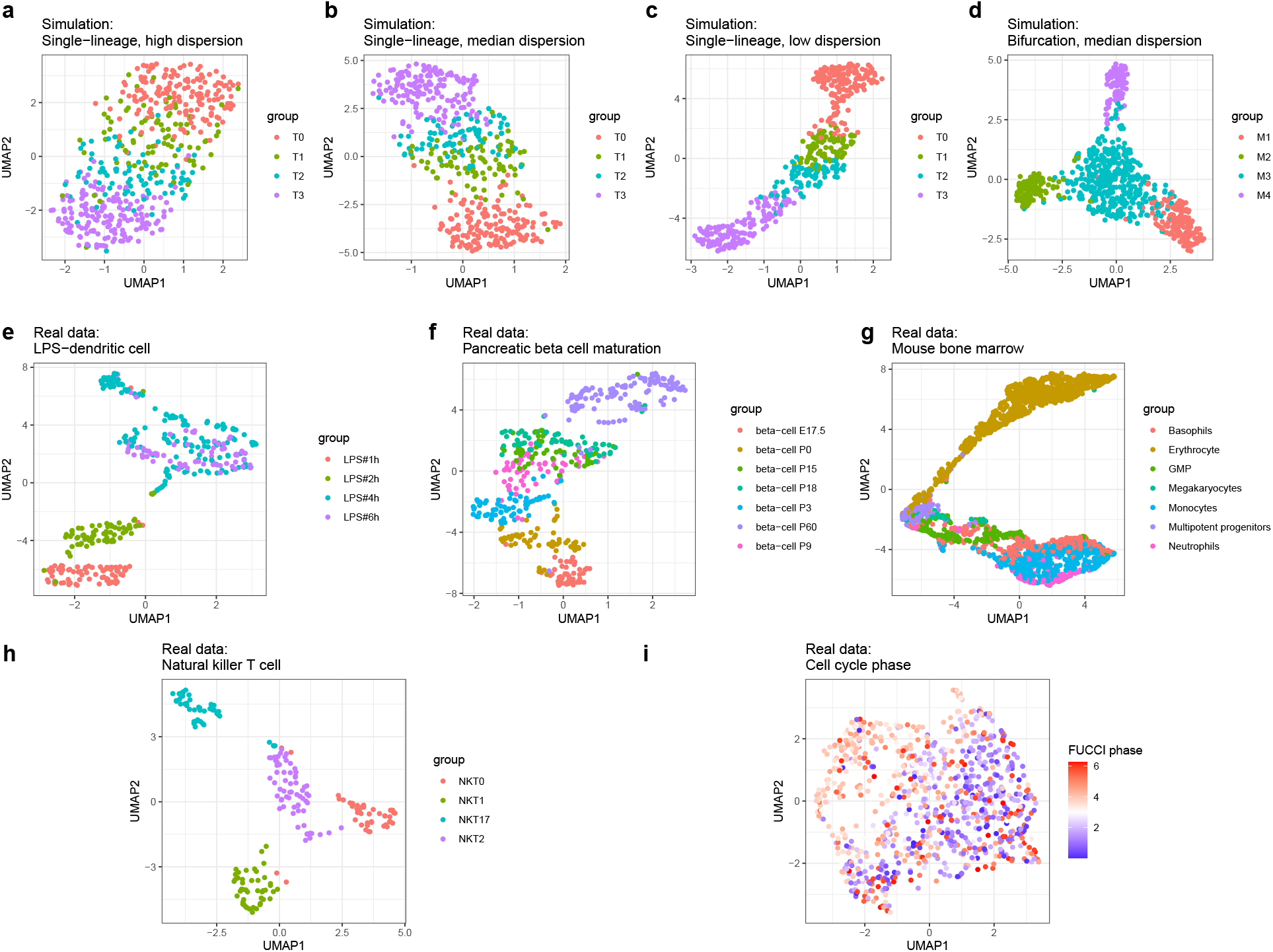
UMAP visualization of datasets. UMAP visualization of all synthetic and real datasets used in this paper. Panels (a)–(d) are synthetic datasets, where the groups correspond to time points. Panels (e)–(h) are real datasets, where the groups correspond to time points in (e) & (f) or annotated cell types in (g) & (h). Panel (i) is a real dataset with external cell cycle information, where the FUCCI phase indicates experimentally measured cell cycle phase.

**Figure S3:**
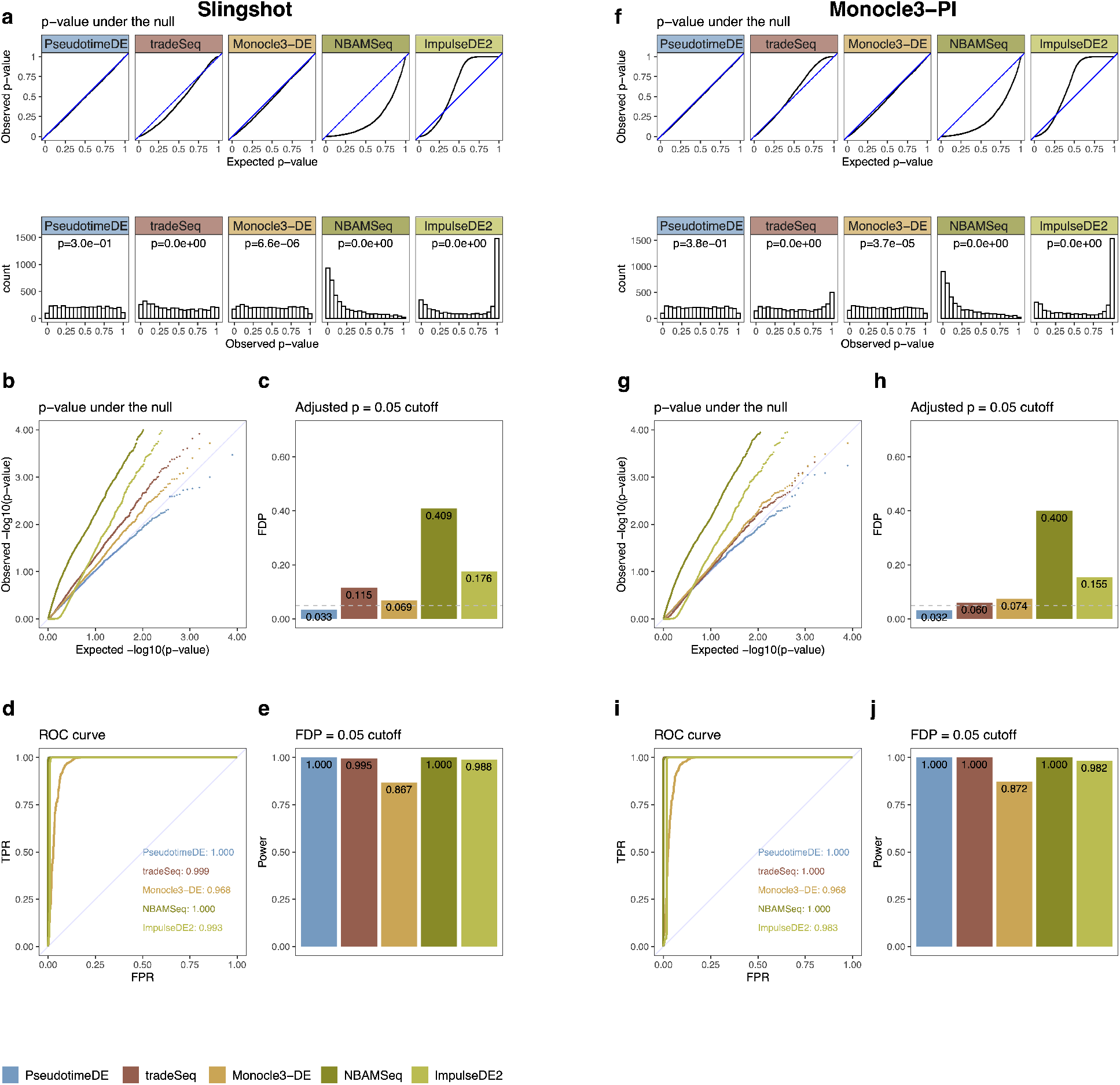
Comparison of five methods (PseudotimeDE, tradeSeq, Monocle3-DE, NBAMSeq, ImpulseDE2) for identifying DE genes along cell pseudotime on synthetic single-lineage data with low dispersion. Left panels (a)–(e) are based on pseudotime inferred by Slingshot; right panels (f)–(j) are based on pseudotime inferred by Monocle3-PI. **(a) & (f)** Distributions of non-DE genes’ observed *p*-values by five DE methods with inferred pseudotime. Top: quantile-quantile plots that compare the empirical quantiles of the observed *p*-values against the expected quantiles of the Uniform[0, 1] distribution. Bottom: histograms of the observed *p*-values. The *p*-values shown on top of histograms are from the Kolmogorov–Smirnov test under the null hypothesis that the distribution is Uniform[0, 1]. The larger the *p*-value, the more uniform the distribution is. Among the five DE methods, PseudotimeDE’s observed *p*-values follow most closely the expected Uniform[0, 1] distribution. **(b) & (g)** Quantile-quantile plots of the same *p*-values as in (a) and (f) on the negative log_10_ scale. PseudotimeDE returns better-calibrated small *p*-values than the other four methods do. **(c) & (h)** FDPs of the five DE methods with the target FDR 0.05 (BH adjusted-*p* ≤ 0.05). PseudotimeDE yields the FDP below 0.05 while other methods do not. **(d) & (i)** ROC curves and AUROC values of the five DE methods. Since the dispersion of data is extremely low, all methods achieves high AUROC. **(e) & (j)** Power of the five DE methods under the FDP = 0.05 cutoff. Due to the same low dispersion reason, all methods achieves high power.

**Figure S4:**
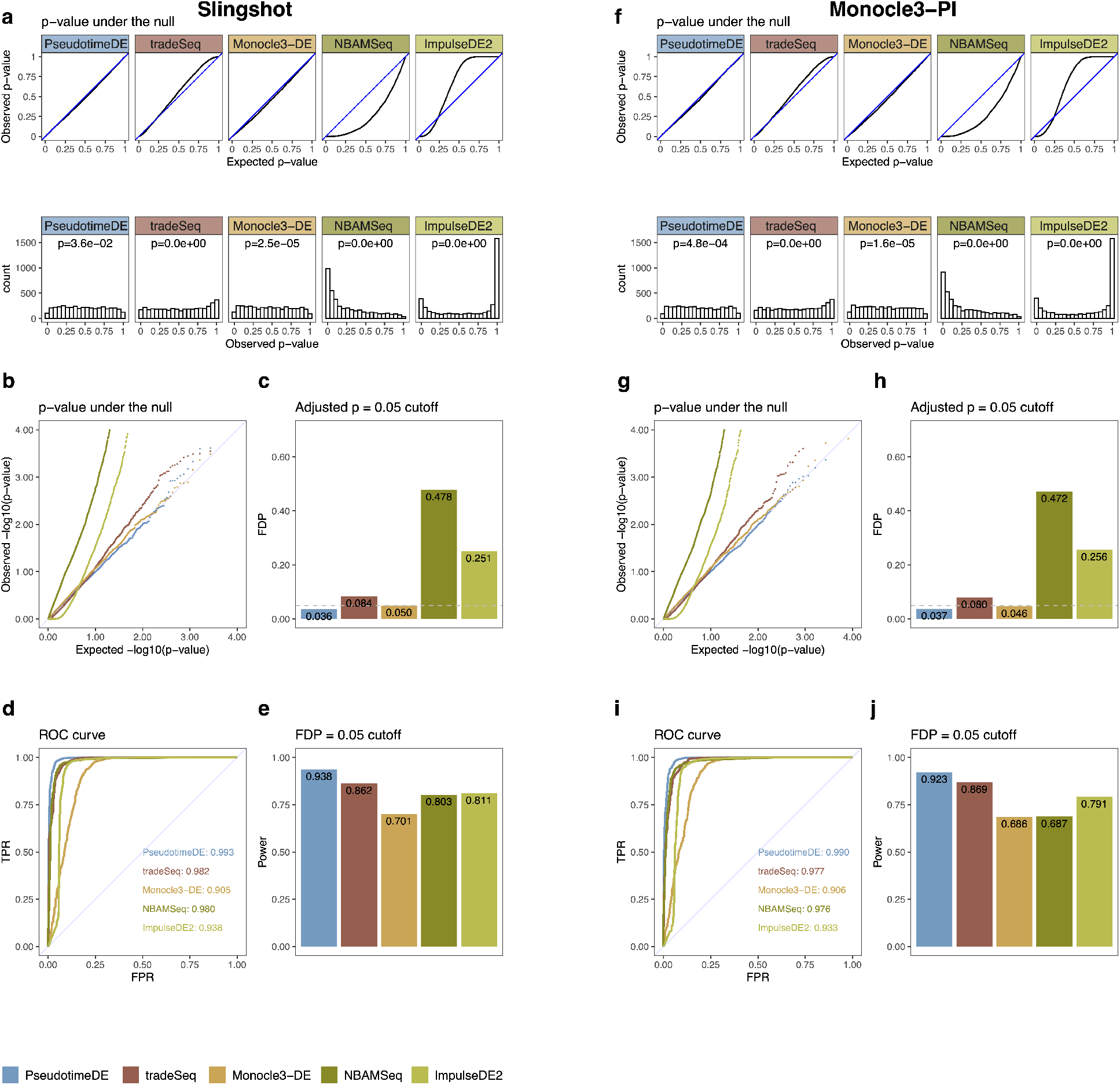
Comparison of five methods (PseudotimeDE, tradeSeq, Monocle3-DE, NBAMSeq, ImpulseDE2) for identifying DE genes along cell pseudotime on synthetic single-lineage data with median dispersion. Left panels (a)–(e) are based on pseudotime inferred by Slingshot; right panels (f)–(j) are based on pseudotime inferred by Monocle3-PI. **(a) & (f)** Distributions of non-DE genes’ observed *p*-values by five DE methods with inferred pseudotime. Top: quantile-quantile plots that compare the empirical quantiles of the observed *p*-values against the expected quantiles of the Uniform[0, 1] distribution. Bottom: histograms of the observed *p*-values. The *p*-values shown on top of histograms are from the Kolmogorov–Smirnov test under the null hypothesis that the distribution is Uniform[0, 1]. The larger the *p*-value, the more uniform the distribution is. Among the five DE methods, PseudotimeDE’s observed *p*-values follow most closely the expected Uniform[0, 1] distribution. **(b) & (g)** Quantile-quantile plots of the same *p*-values as in (a) and (f) on the negative log_10_ scale. PseudotimeDE returns better-calibrated small *p*-values than the other four methods do. **(c) & (h)** FDPs of the five DE methods with the target FDR 0.05 (BH adjusted-*p* ≤ 0.05).PseudotimeDE yields the FDP below 0.05 while other methods do not. **(d) & (i)** ROC curves and AUROC values of the five DE methods. PseudotimeDE achieves the highest AUROC. **(e) & (j)** Power of the five DE methods under the FDP = 0.05 cutoff. PseudotimeDE achieves the highest power.

**Figure S5:**
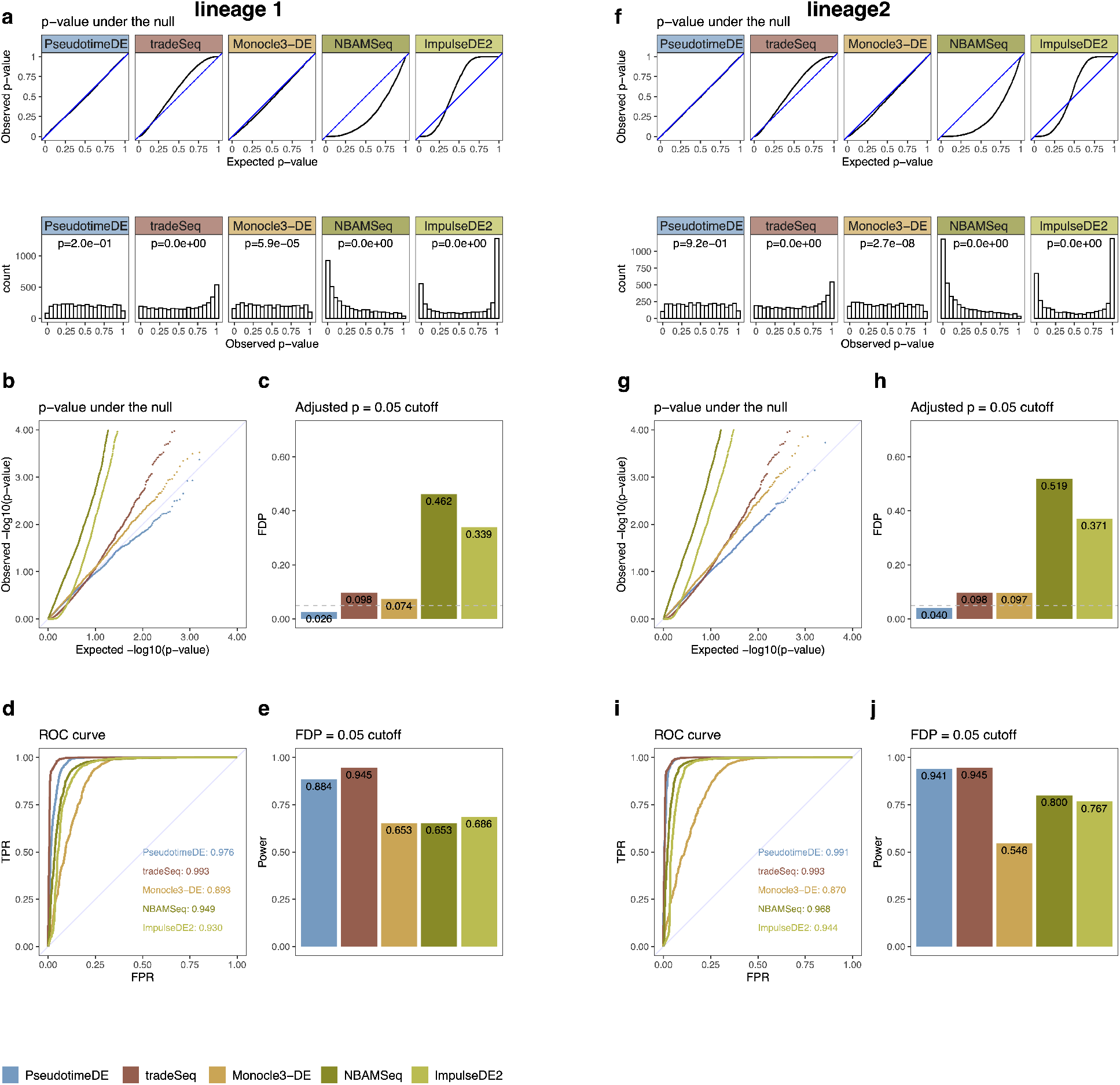
Comparison of five methods (PseudotimeDE, tradeSeq, Monocle3-DE, NBAMSeq, ImpulseDE2) for identifying DE genes along cell pseudotime on synthetic bifurcation data. Pseudotime is inferred by Slingshot. Left panels (a)–(e) are based on lineage 1 of two lineages in bifurcation data; right panels (f)–(j) are based on lineage 2. **(a) & (f)** Distributions of non-DE genes’ observed *p*-values by five DE methods with inferred pseudotime. Top: quantile-quantile plots that compare the empirical quantiles of the observed *p*-values against the expected quantiles of the Uniform[0, 1] distribution. Bottom: histograms of the observed *p*-values. The *p*-values shown on top of histograms are from the Kolmogorov–Smirnov test under the null hypothesis that the distribution is Uniform[0, 1]. The larger the *p*-value, the more uniform the distribution is. Among the five DE methods, PseudotimeDE’s observed *p*-values follow most closely the expected Uniform[0, 1] distribution. **(b) & (g)** Quantile-quantile plots of the same *p*-values as in (a) and (f) on the negative log_10_ scale. PseudotimeDE returns better-calibrated small *p*-values than the other four methods do. **(c) & (h)** FDPs of the five DE methods with the target FDR 0.05 (BH adjusted-*p* ≤ 0.05). PseudotimeDE yields the FDP below 0.05 while other methods do not. **(d) & (i)** ROC curves and AUROC values of the five DE methods. PseudotimeDE achieves the second highest AUROC which is close to the highest value. **(e) & (j)** Power of the five DE methods under the FDP = 0.05 cutoff. PseudotimeDE achieves the second highest power which is close to the highest value.

**Figure S6:**
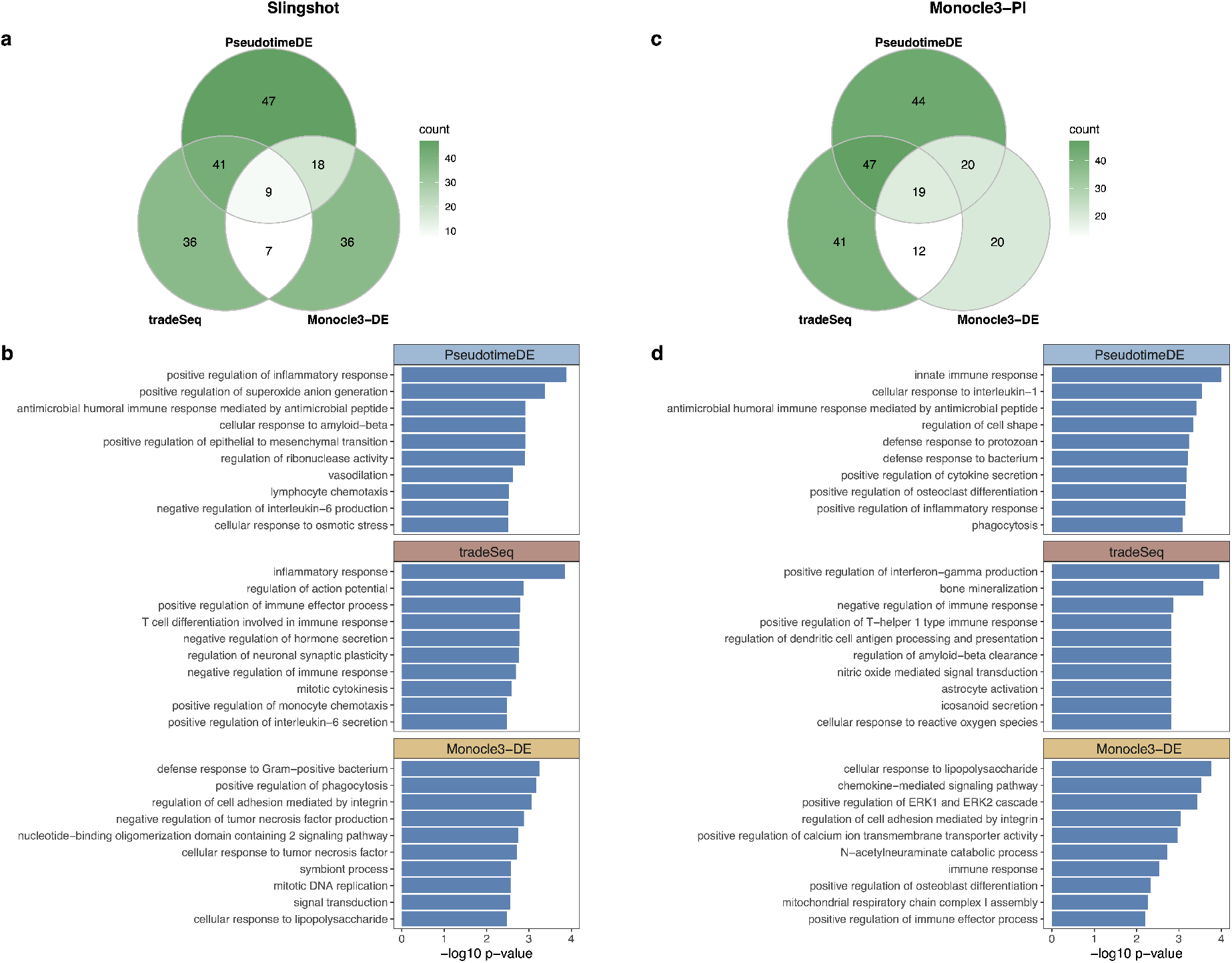
GO analysis of DE genes identified in the LPS-dendritic cell dataset. Left panels (a)–(b) are based on pseudotime inferred by Slingshot; right panels (c)–(d) are based on pseudotime inferred by Monocle3-PI. **(a) & (c)** Numbers of GO terms enriched (*p* < 0.01) in the significant DE genes found by each method in Fig. 4 (b) & (f). **(b) & (d)** Top 10 enriched GO terms for each DE method.

**Figure S7:**
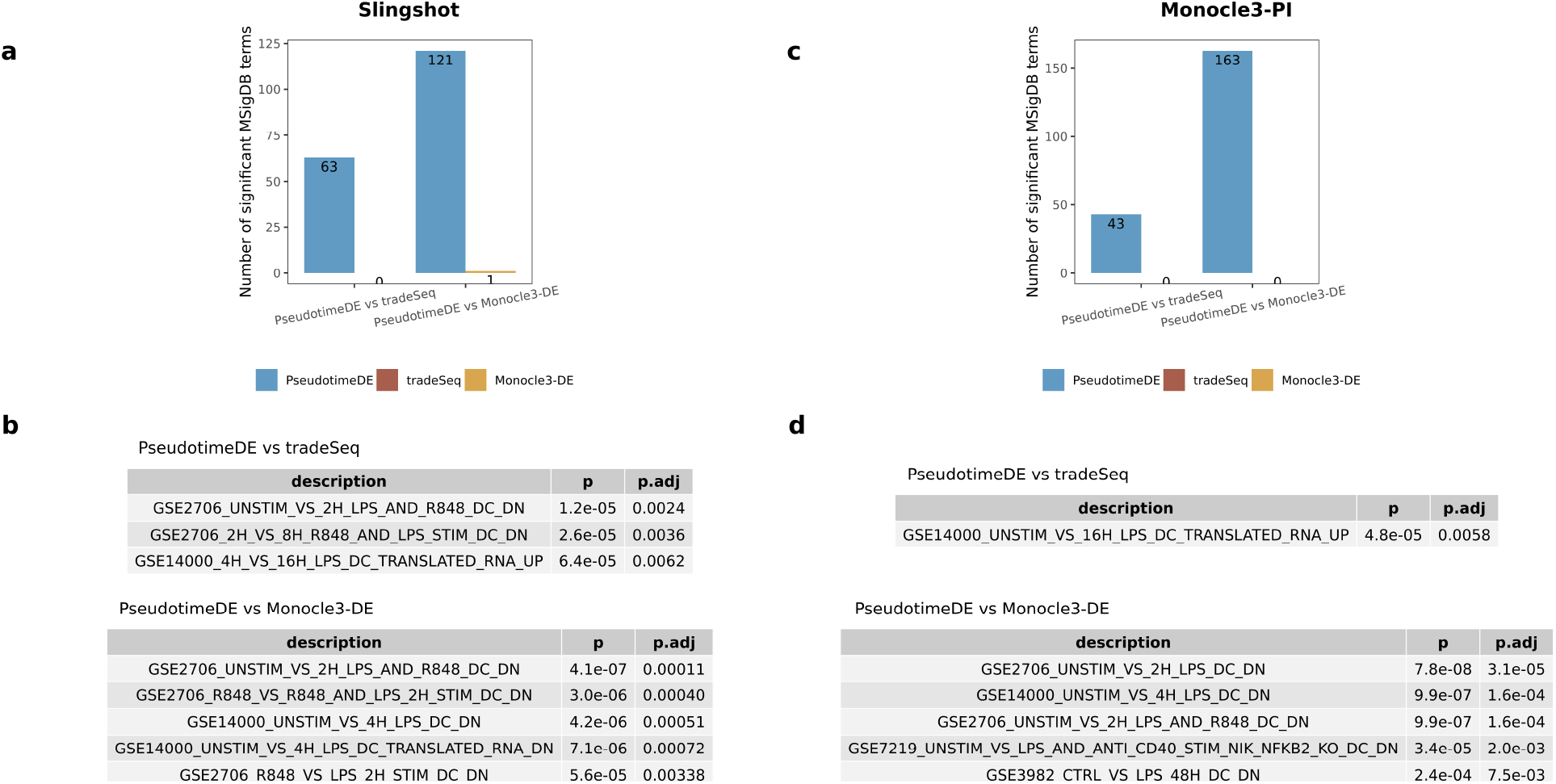
MSigDB over-representation analysis of DE genes identified in the LPS-dendritic cell dataset. Left panels (a)–(b) are based on pseudotime inferred by Slingshot; right panels (c)–(d) are based on pseudotime inferred by Monocle3-PI. **(a) & (c)** Numbers of MSigDB terms enriched (BH adjusted-*p* < 0.01) in the significant DE genes specifically found by PseudotimeDE or tradeSeq/Monocle3-DE in pairwise comparisons between PseudotimeDE and tradeSeq/Monocle3-DE in Fig. 4 (b) & (f). **(b) & (d)** Example MSigDB terms enriched in the Pseudotime-specific DE genes in (a) & (b). The explanation of terms can be found from MSigDB. All listed terms are related to the response of dendritic cells (DC) stimulated by LPS.

**Figure S8:**
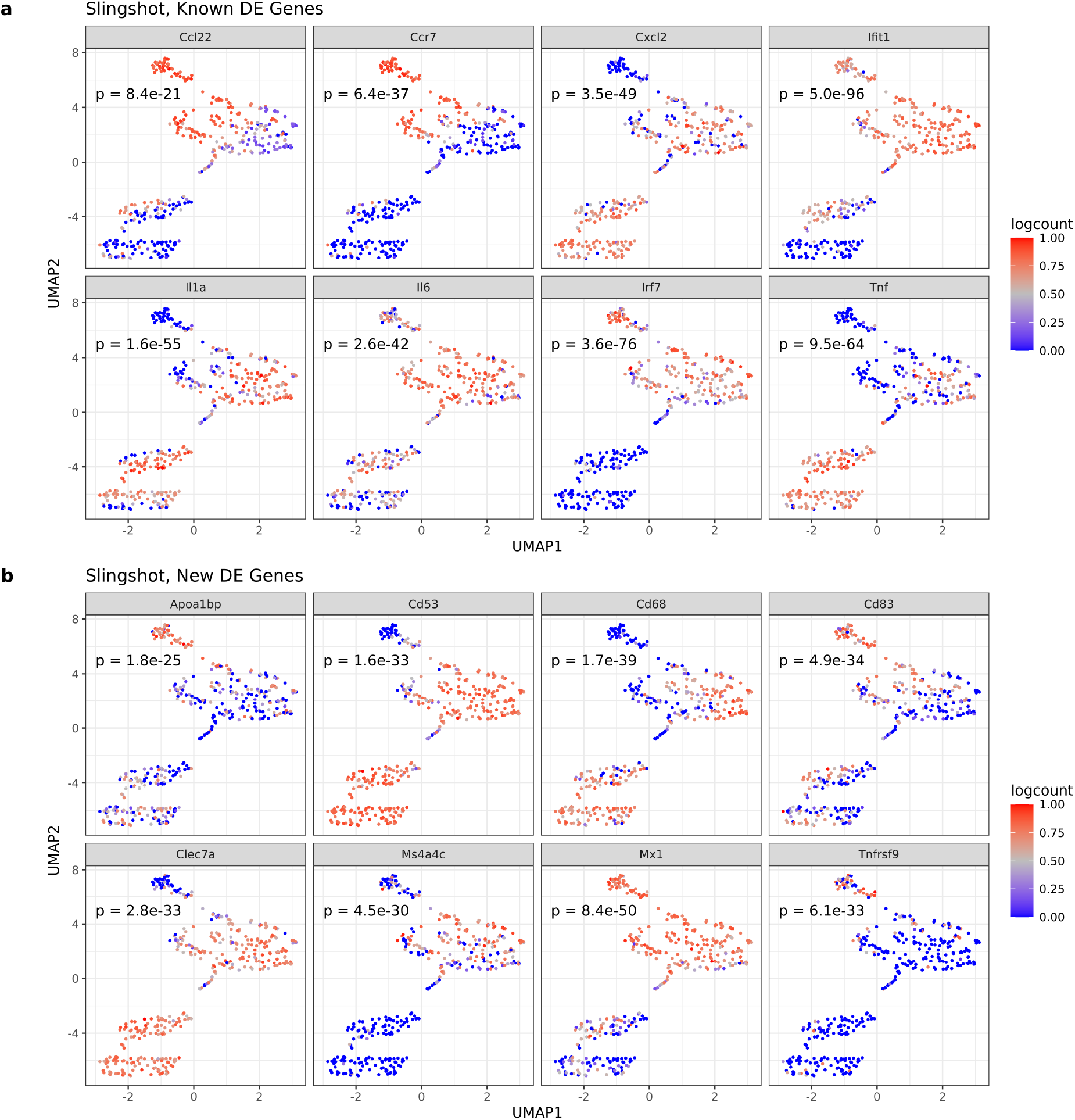
UMAP visualization of example DE genes identified by PseudotimeDE, using Slingshot as the pseudotime inference method, in the LPS-dendritic cell dataset. *p*-values returned by PseudotimeDE are reported for all the 16 example genes. **(a)** Examples of known DE genes, which have been reported as highly confident DE genes in the original study [31]. **(b)** Examples of new DE genes, which are identified by PseudotimeDE but found as non-DE by either tradeSeq or Monocle3-DE.

**Figure S9:**
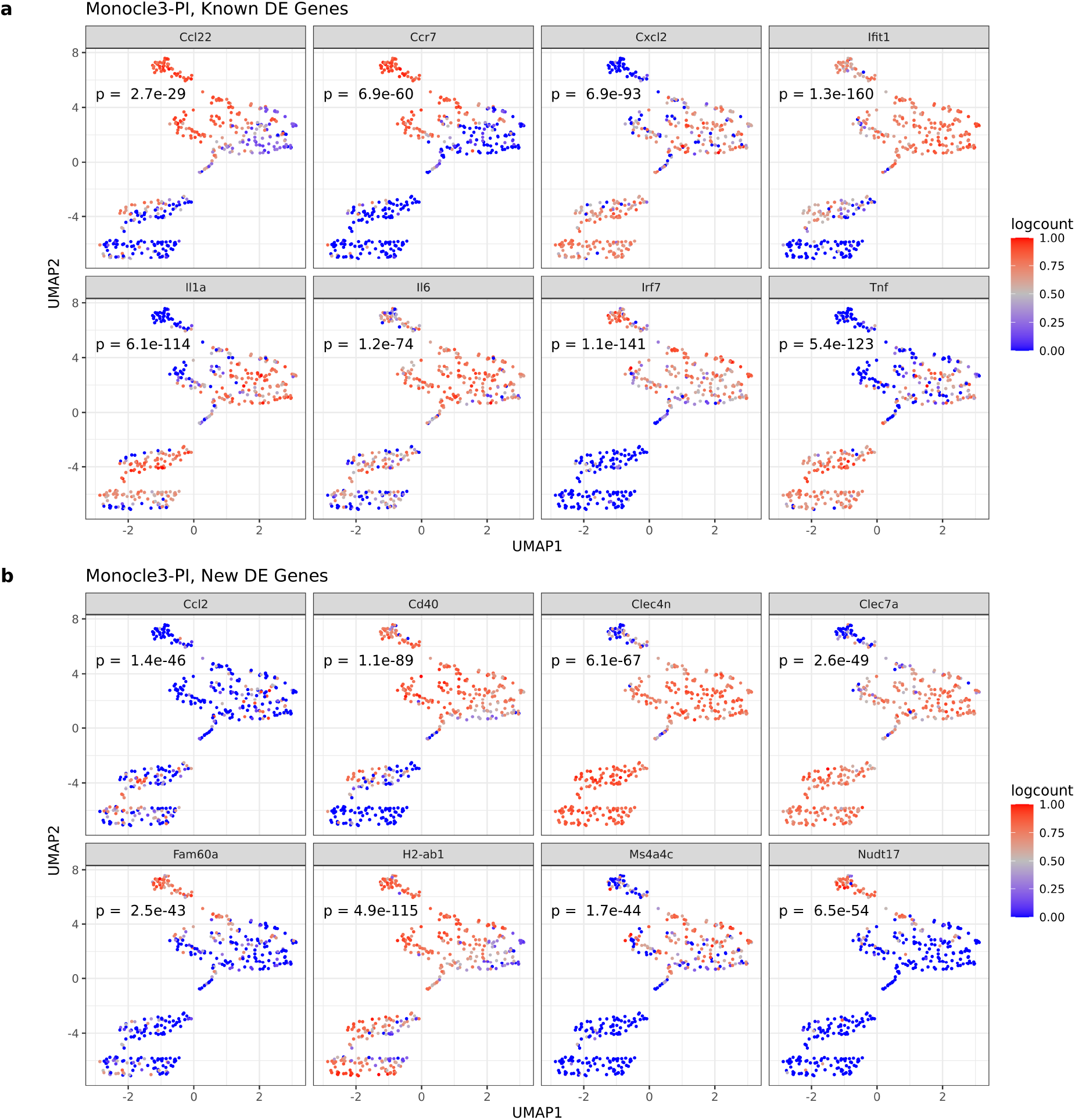
UMAP visualization of example DE genes identified by PseudotimeDE, using Monocle3-PI as the pseudotime inference method, in the LPS-dendritic cell dataset. *p*-values returned by PseudotimeDE are reported for all the 16 example genes. **(a)** Examples of known DE genes, which have been reported as highly confident DE genes in the original study [31]. **(b)** Examples of new DE genes, which are identified by PseudotimeDE but found as non-DE by either tradeSeq or Monocle3-DE.

**Figure S10:**
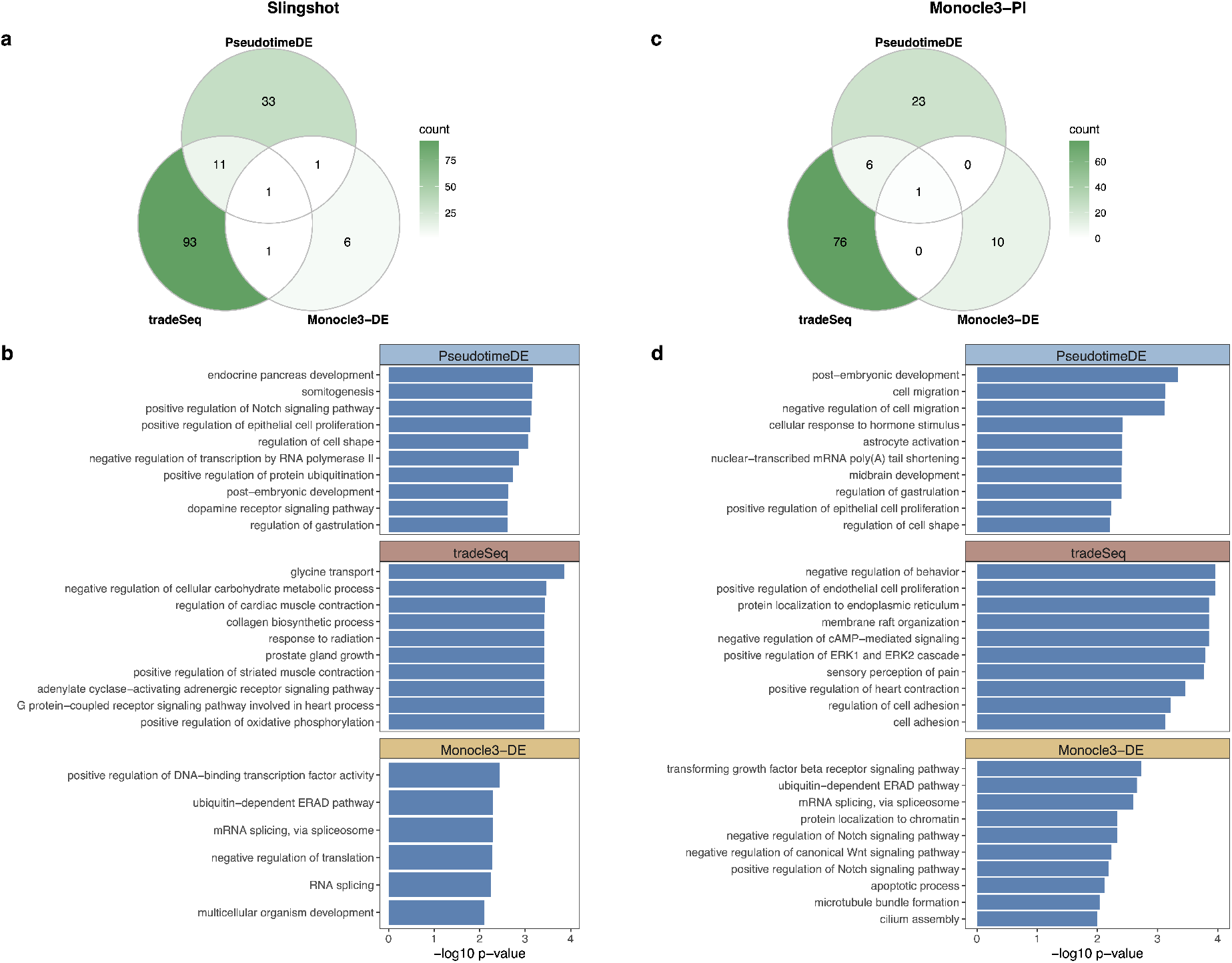
GO analysis of DE genes identified in the pancreatic beta cell maturation dataset. Left panels (a)–(b) are based on pseudotime inferred by Slingshot; right panels (c)–(d) are based on pseudotime inferred by Monocle3-PI. **(a) & (c)** Numbers of GO terms enriched (*p* < 0.01) in the significant DE genes found by each method in Fig. 5 (b) & (f). **(b) & (d)** Top 10 significant GO terms from each DE method.

**Figure S11:**
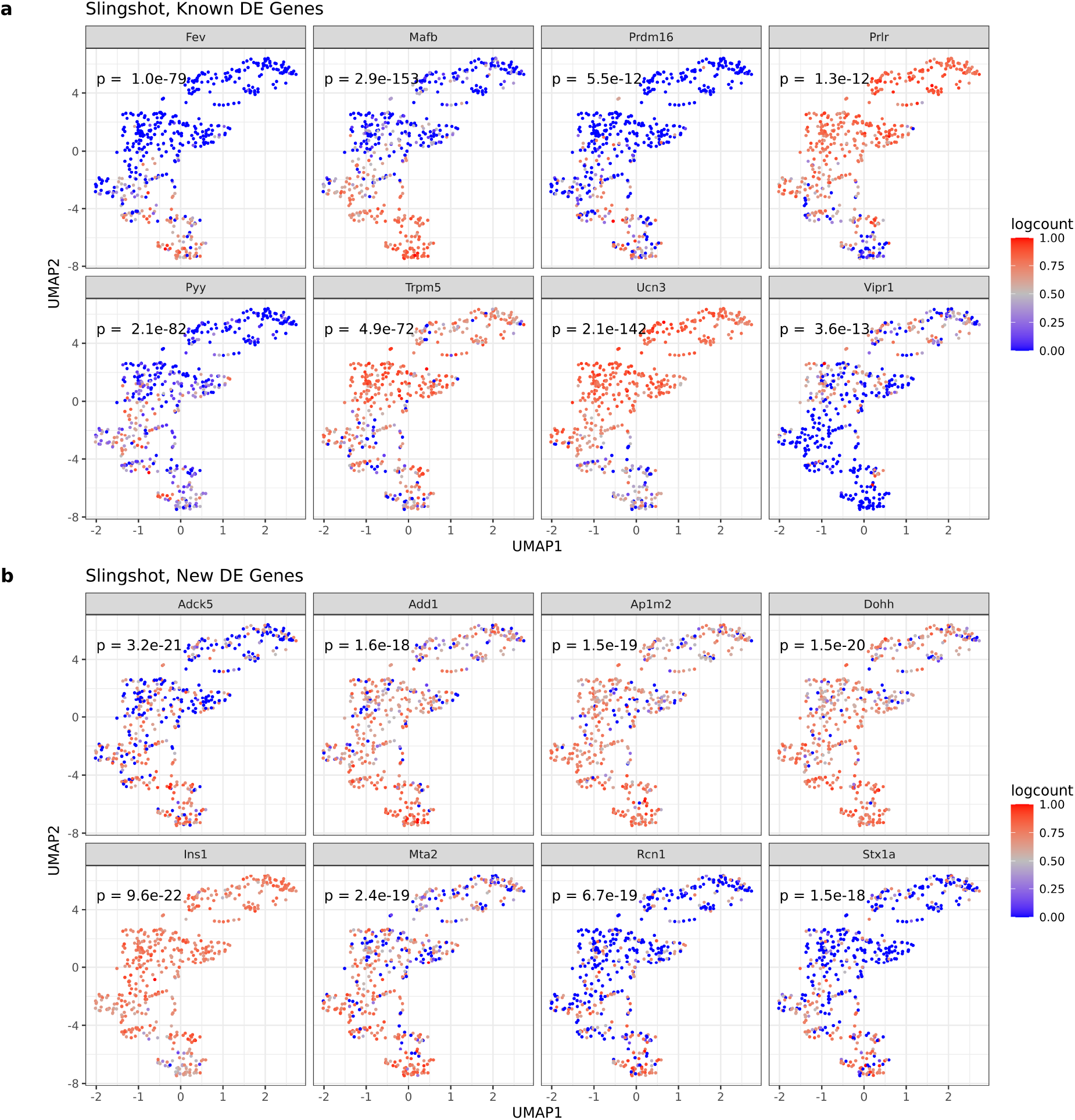
UMAP visualization of example DE genes identified by PseudotimeDE, using Slingshot as the pseudotime inference method, in the pancreatic beta cell maturation cell dataset. *p*-values returned by PseudotimeDE are reported for all the 16 example genes. **(a)** Examples of known DE genes, which have been reported as highly confident DE genes in the original study [33]. **(b)** Examples of new DE genes, which are identified by PseudotimeDE but found as non-DE by either tradeSeq or Monocle3-DE.

**Figure S12:**
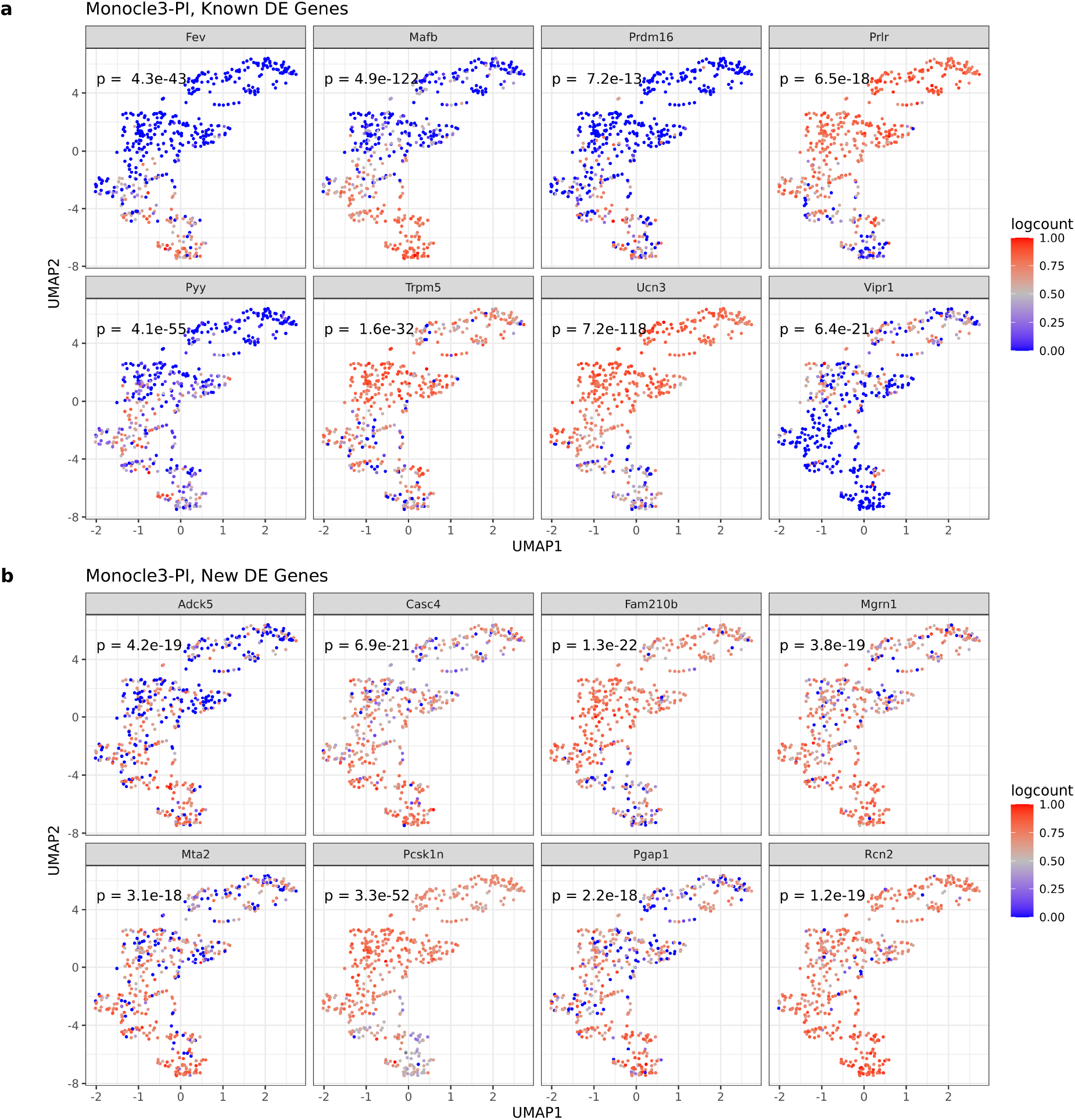
UMAP visualization of example DE genes identified by PseudotimeDE, using Monocle3-PI as the pseudotime inference method, in the pancreatic beta cell maturation cell dataset. *p*-values returned by PseudotimeDE are reported for all the 16 example genes. **(a)** Examples of known DE genes, which have been reported as highly confident DE genes in the original study [33]. **(b)** Examples of new DE genes, which are identified by PseudotimeDE but found as non-DE by either tradeSeq or Monocle3-DE.

**Figure S13:**
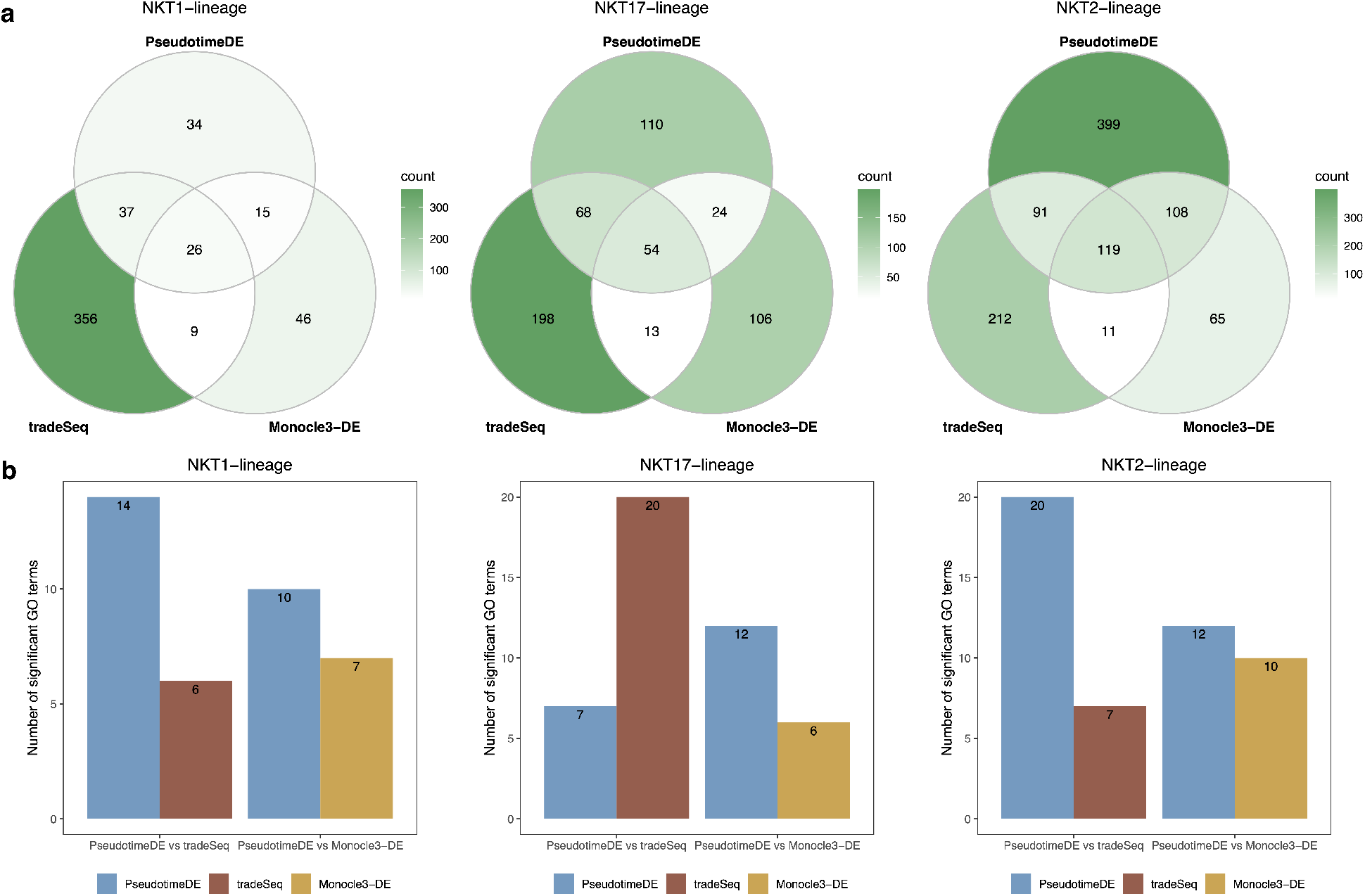
GO analysis of DE genes identified in the natural killer T cell dataset. **(a)** Venn plots showing the overlaps of the significant DE genes (BH adjusted-*p* ≤ 0.05) identified by the three DE methods. **(b)** Numbers of GO terms enriched (*p* < 0.05) in the significant DE genes specifically found by PseudotimeDE or tradeSeq/Monocle3-DE in pairwise comparisons between PseudotimeDE and tradeSeq/Monocle3-DE in (a).

**Figure S14:**
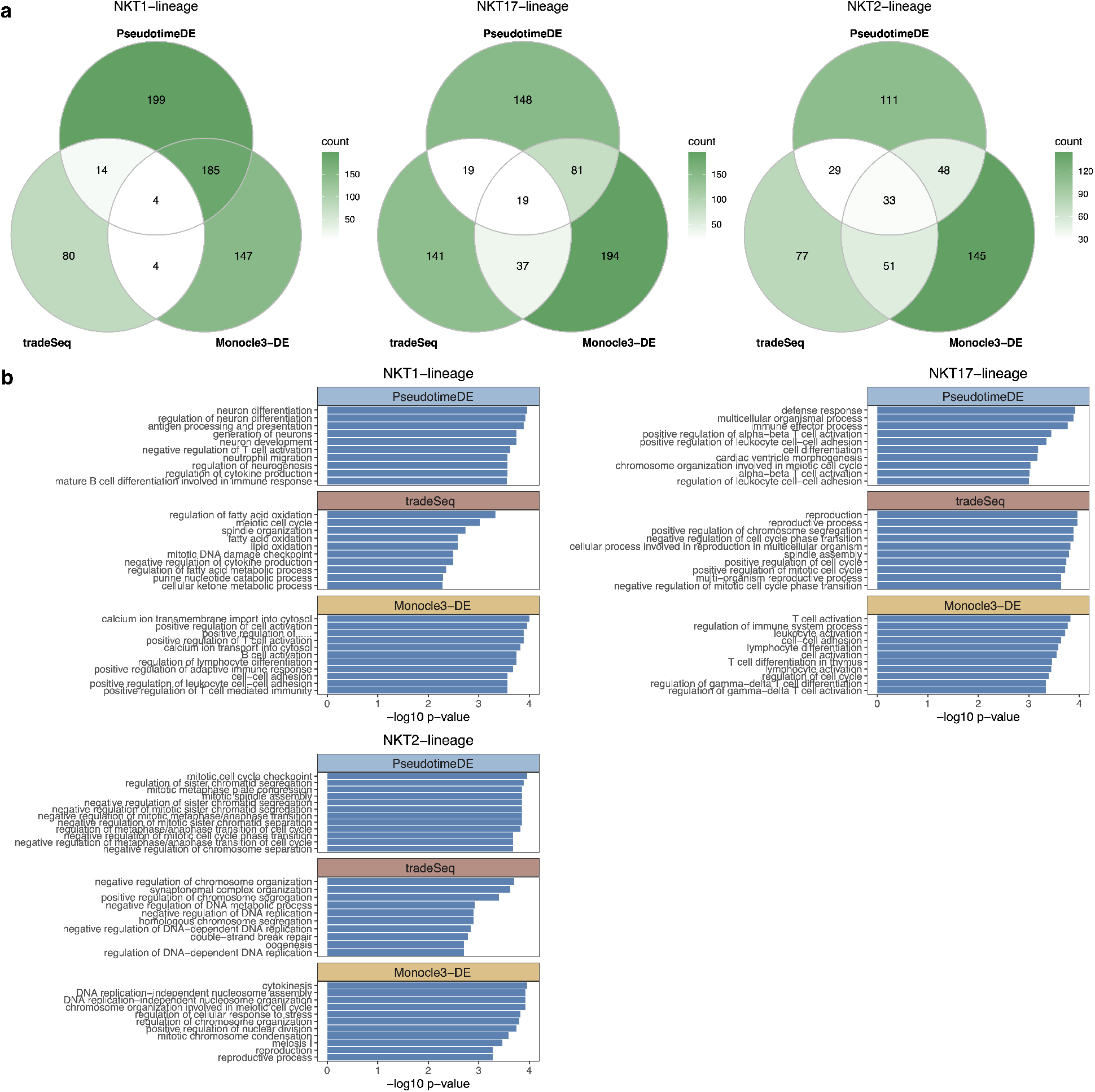
GO analysis of DE genes identified in the natural killer T cell dataset. **(a)** Numbers of GO terms enriched (*p* < 0.05) in the significant DE genes found by each method. **(b)** Top 10 enriched GO terms for each DE method.

**Figure S15:**
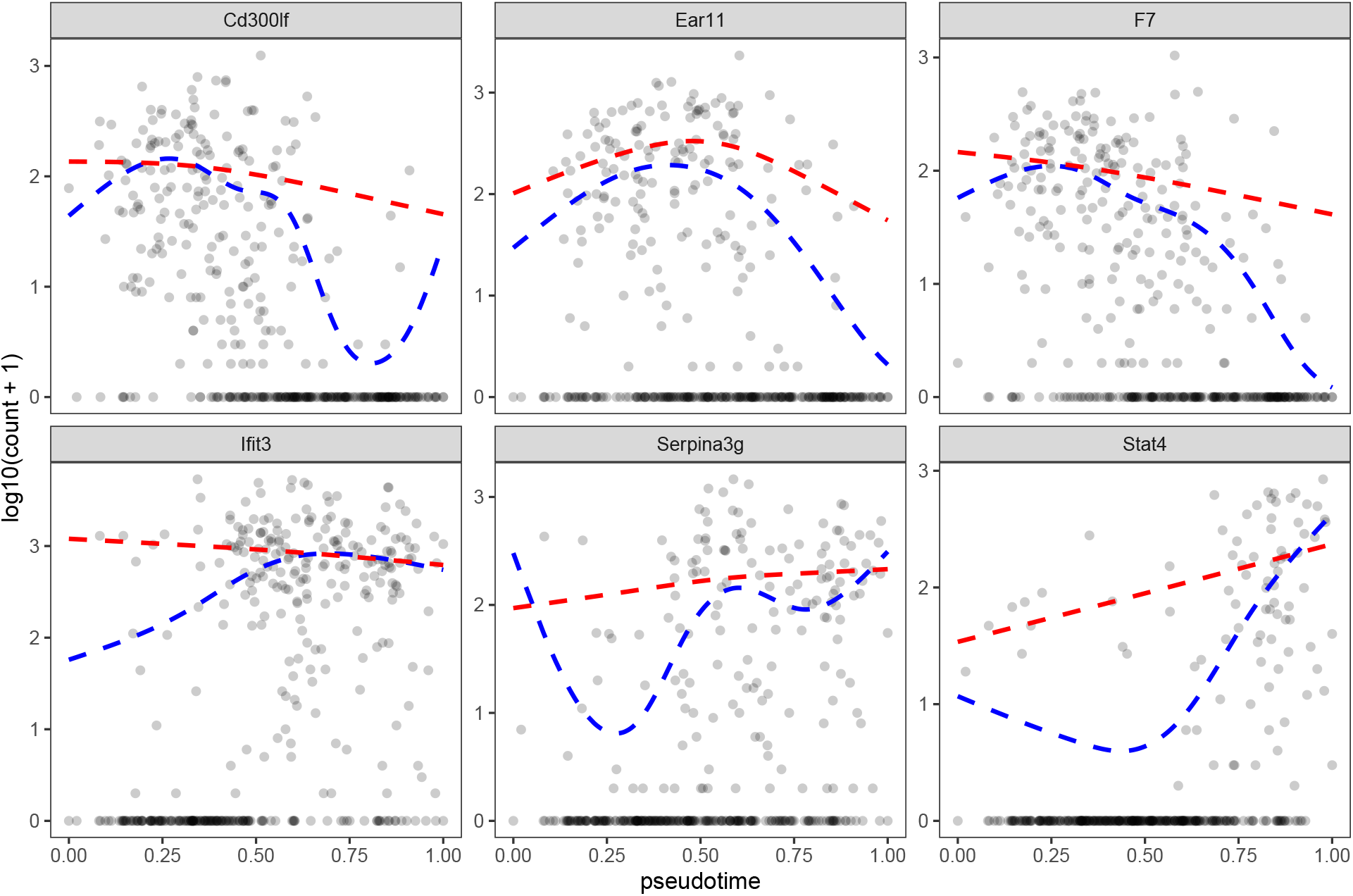
Comparison of NB-GAM and ZINB-GAM on the LPS-dendritic cell dataset with Slingshot pseudotime. Example fitted results of NB-GAM / ZINB-GAM on six genes from the LPS-dendritic cell dataset with pseudotime inferred by Slingshot. NB-GAM yields small *p*-values (*p* < 1*e* − 10) and ZINB-GAM yields large *p*-values (*p* > 0.01). Dashed blue lines and red lines are the fitted curves by NB-GAM and ZINB-GAM, respectively.

**Figure S16:**
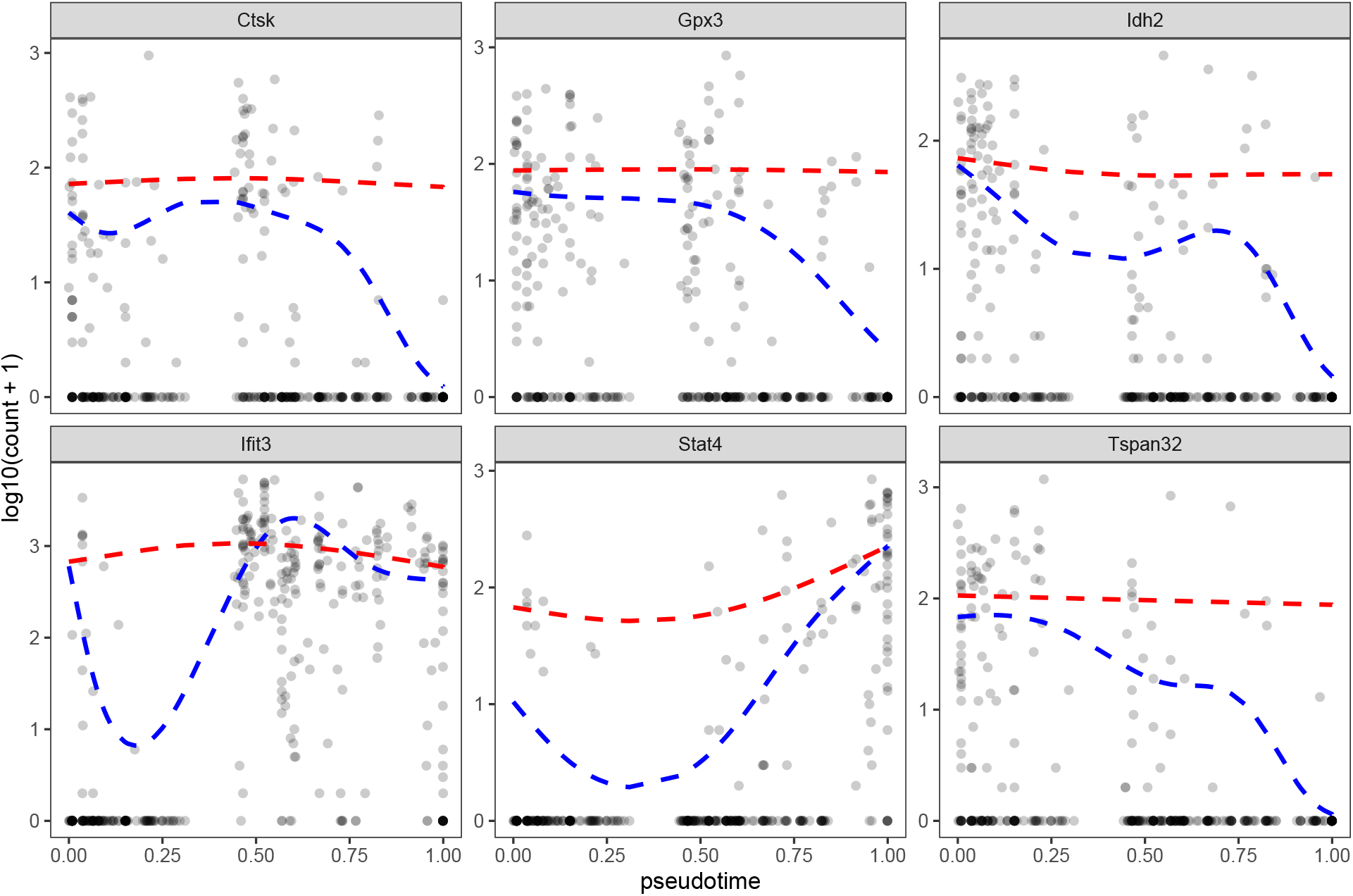
Comparison of NB-GAM and ZINB-GAM on the LPS-dendritic cell dataset with Monocle3-PI pseudotime. Example fitted results of NB-GAM / ZINB-GAM on six genes from the LPS-dendritic cell dataset with pseudotime inferred by Monocle3-PI. NB-GAM yields small *p*-values (*p* < 1*e* − 10) and ZINB-GAM yields large *p*-values (*p* > 0.01). Dashed blue lines and red lines are the fitted curves by NB-GAM and ZINB-GAM, respectively.

**Figure S17:**
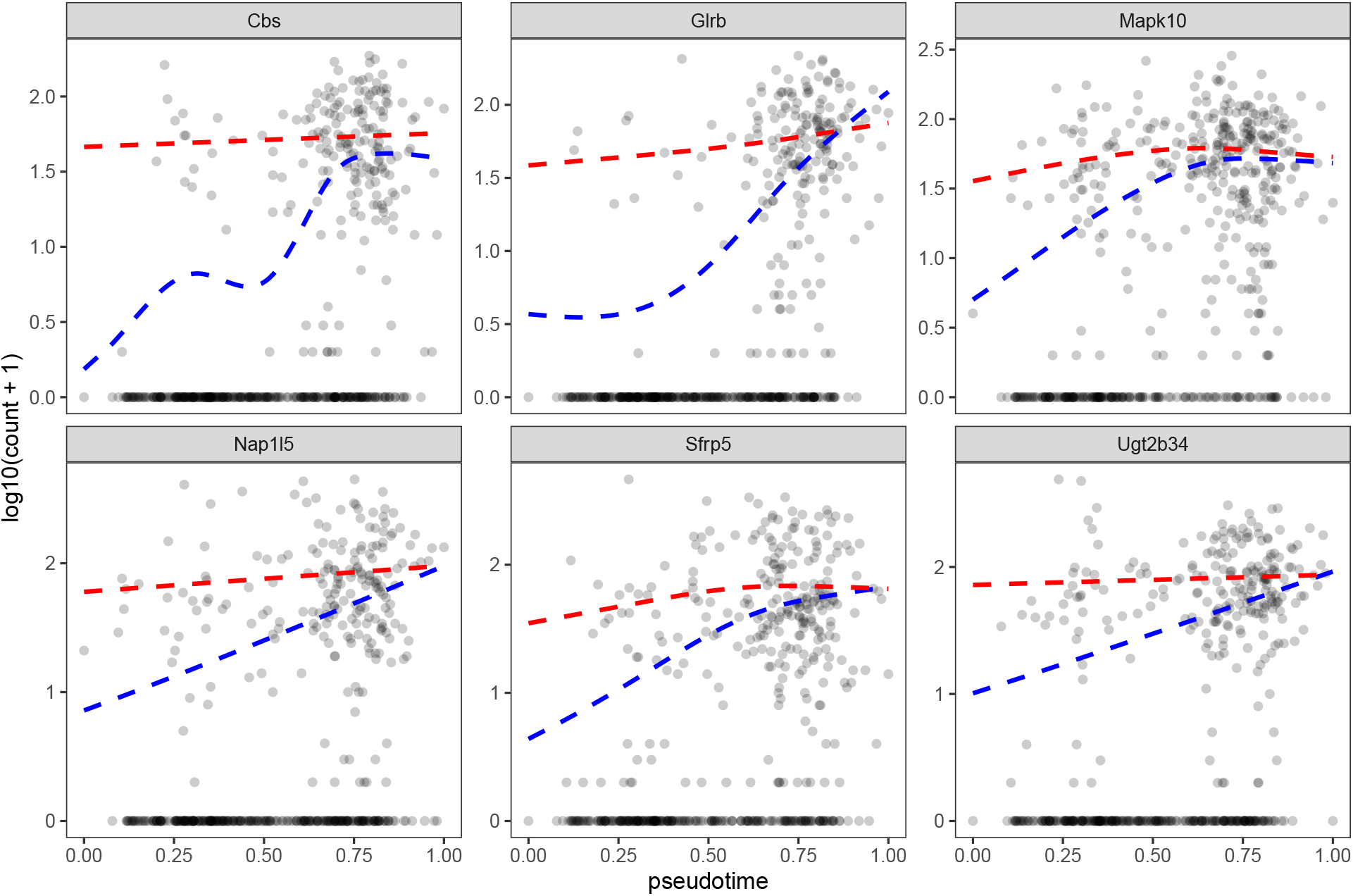
Comparison of NB-GAM and ZINB-GAM on the pancreatic beta cell maturation dataset with Slingshot pseudotime. Example fitted results of NB-GAM / ZINB-GAM on six genes fromthe pancreatic beta cell maturation dataset with pseudotime inferred by Slingshot. NB-GAM yields small *p*-values (*p* < 1*e* − 10) and ZINB-GAM yields large *p*-values (*p* > 0.01). Dashed blue lines and red lines are the fitted curves by NB-GAM and ZINB-GAM, respectively.

**Figure S18:**
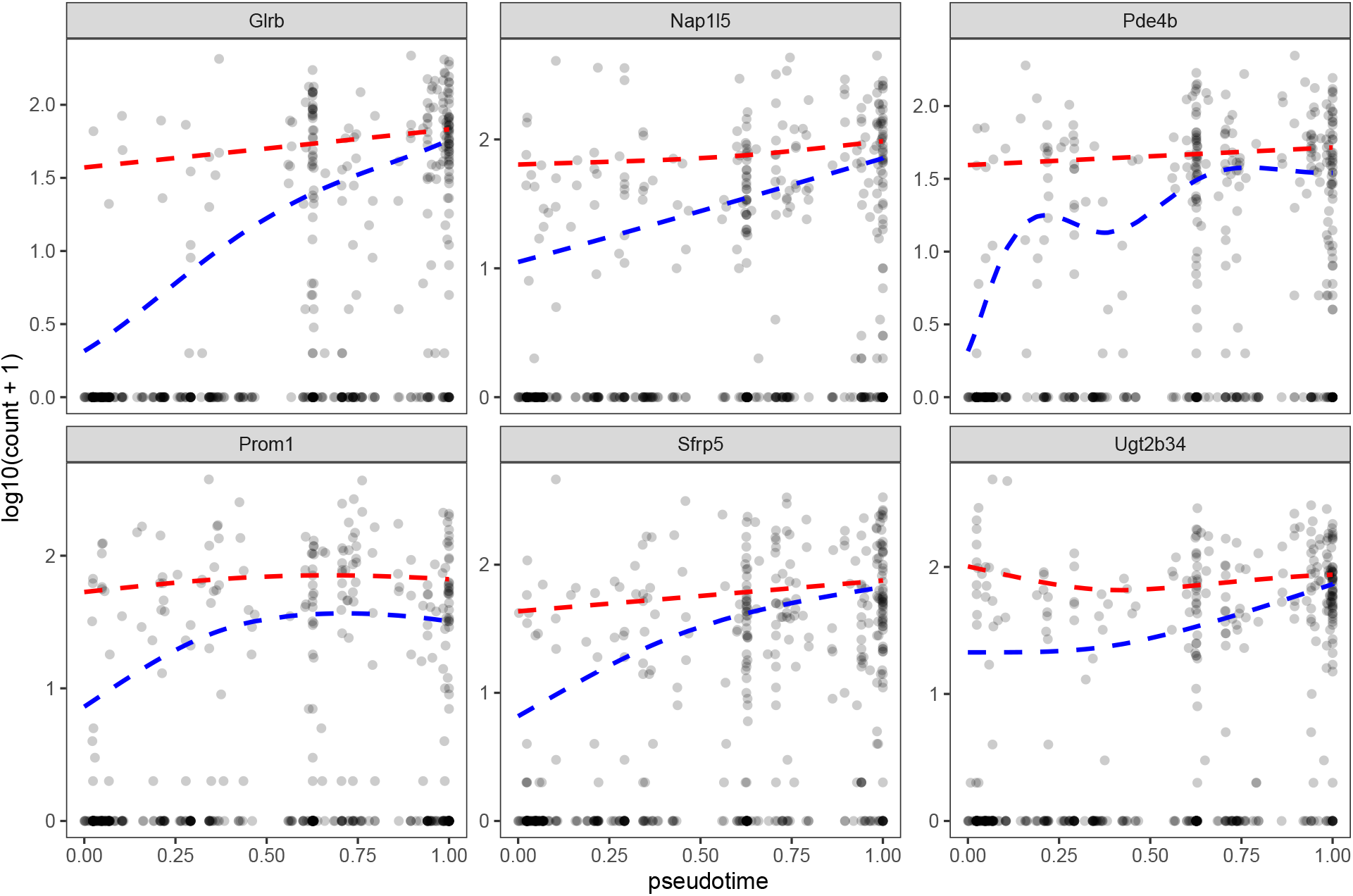
Comparison of NB-GAM and ZINB-GAM on the pancreatic beta cell maturation cell dataset with Slingshot pseudotime. Example fitted results of NB-GAM / ZINB-GAM on six genes from the pancreatic beta cell maturation with pseudotime inferred by Monocle3-PI. NB-GAM yields small *p*-values (*p* < 1*e* − 10) and ZINB-GAM yields large *p*-values (*p* > 0.01). Dashed blue lines and red lines are the fitted curves by NB-GAM and ZINB-GAM, respectively.

**Figure S19:**
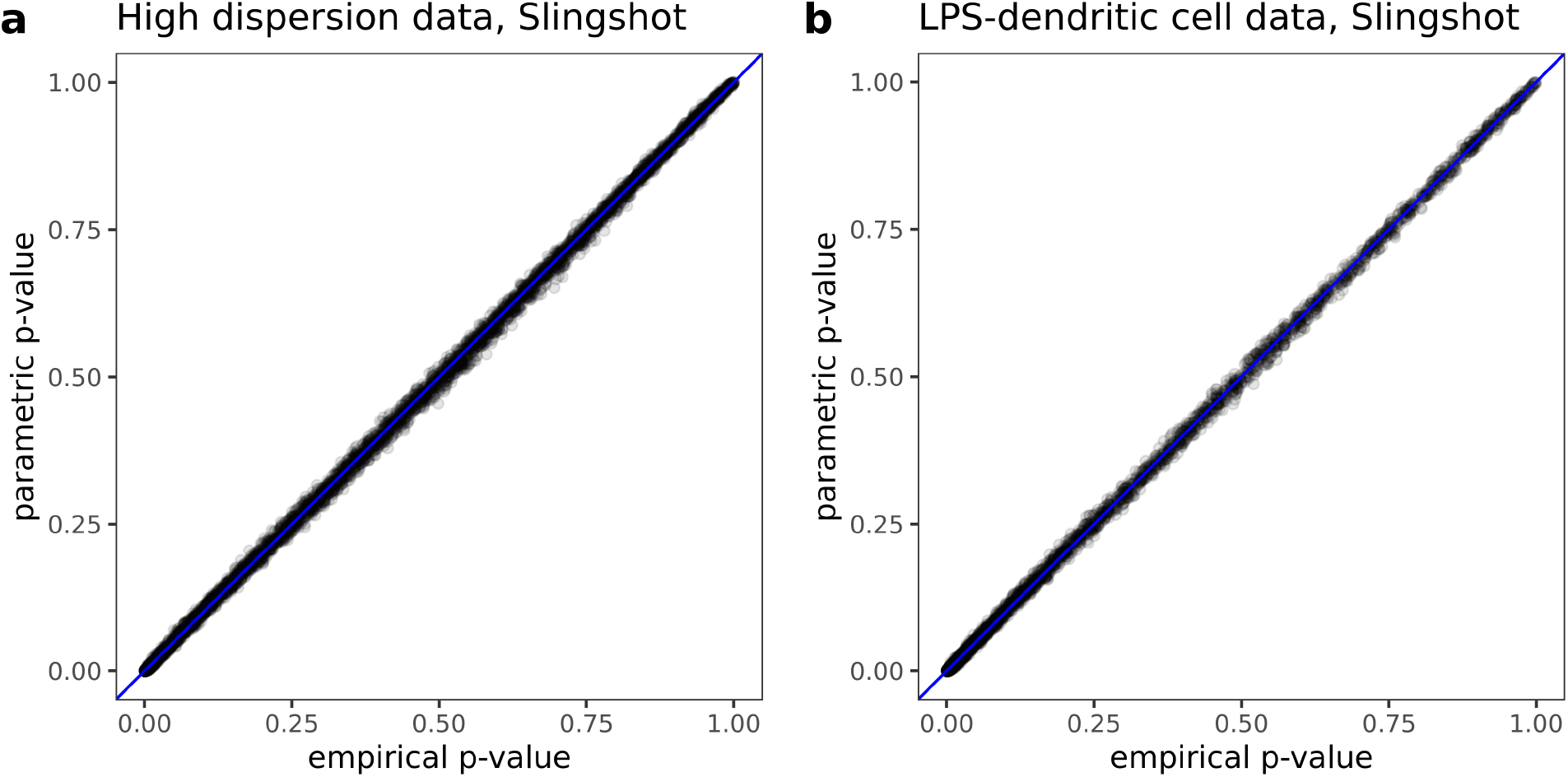
Comparison of empirical *p*-value and parametric *p*-value. Scatter plot of empirical *p*-values and parametric *p*-values (see **Methods**). **(a)** Scatter plot based on synthetic high dispersion dataset and pseudotime inferred by Slingshot. **(b)** Scatter plot based on LPS-dendritic cell dataset and pseudotime inferred by Slingshot. The parametric *p*-values are perfectly correlated with empirical *p*-values, suggesting that the parametric model well captures the estimated null distribution of test statistics.

**Figure S20:**
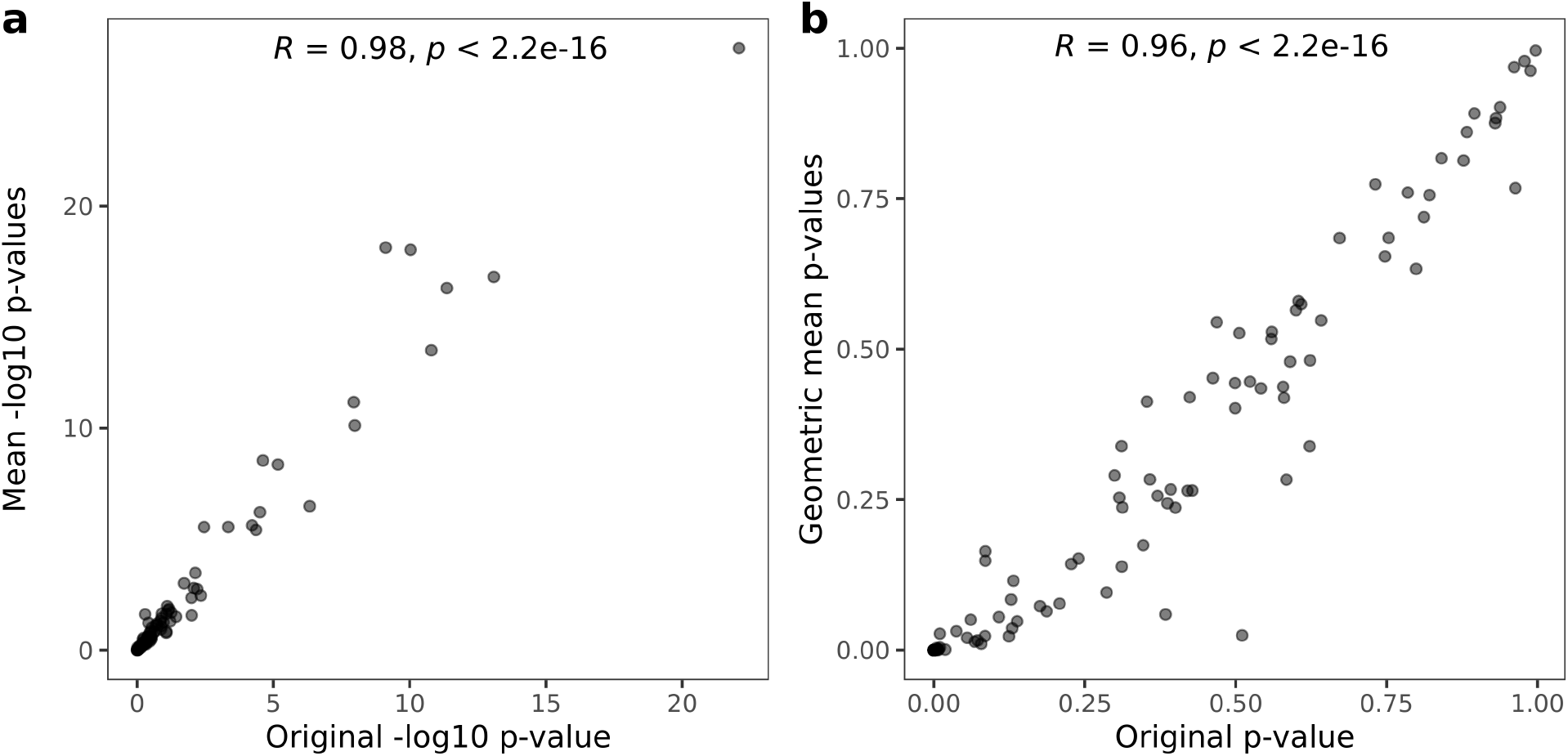
Comparison of *p*-values using 1000 subsamples and *p*-values using 100 subsamples. The *p*-values are based on based on synthetic high dispersion dataset and pseudotime inferred by Slingshot. **(a)** Scatter plot of the original *p*-value using 1000 subsamples on negative – log_10_ scale and the mean of 50 *p*-values using 100 subsamples on negative – log_10_ scale. The strict linearity (Pearson correlation coefficient *R* = 0.98) suggests that using 100 subsamples yield similar *p*-values as using 1000 subsamples on log scale. **(b)** Scatter plot of the original *p*-value using 1000 subsamples and the geometric mean of 50 *p*-values using 100 subsamples. The strict linearity (Pearson correlation coefficient *R* = 0.96) suggests that using 100 subsamples yield similar *p*-values as using 1000 subsamples on raw scale.

**Figure S21:**
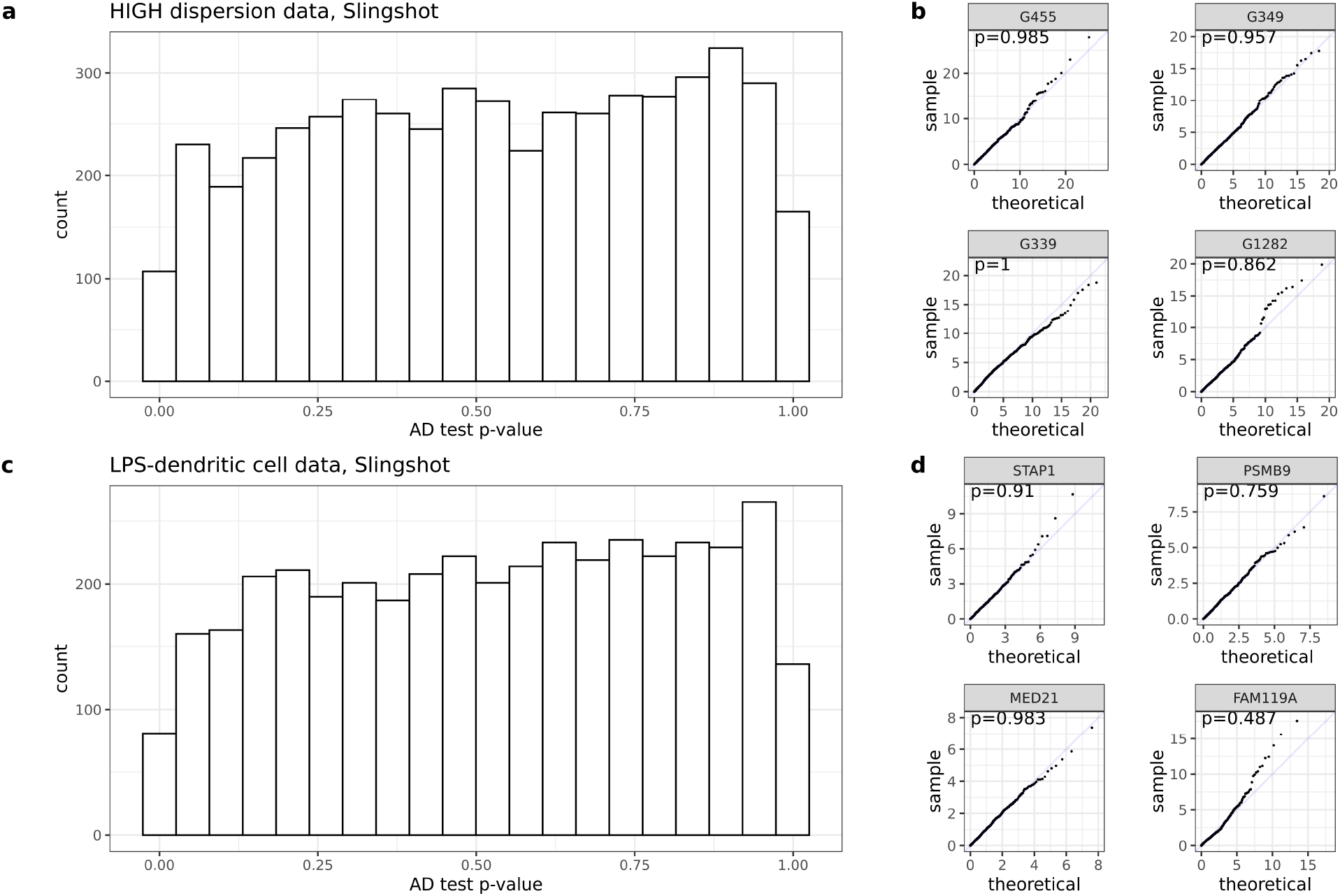
Goodness-of-fit of the parametric distribution. The *p*-values are from Anderson-Darling (AD) test, which measures the goodness-of-fit of the gamma/two-component gamma mixture distribution to the empirical null distribution generated by subsampling and permutation. **(a)** Histogram of AD test *p*-values based on the synthetic high dispersion dataset and pseudotime inferred by Slingshot. The distribution is approximately Uniform[0,1], indicating that the parametric distribution fits the empirical null distribution well. **(b)** Quantile-quantile plots comparing the empirical null distribution and its corresponding parametric fit for four random genes. **(c)** Histogram of AD test *p*-values based on the LPS-dendritic cell dataset and pseudotime inferred by Slingshot. The distribution is approximately Uniform[0,1], indicating that the parametric distribution fits the empirical null distribution well. **(d)** Quantile-quantile plots comparing the empirical null distribution and its corresponding parametric fit for four random genes.

**Figure S22:**
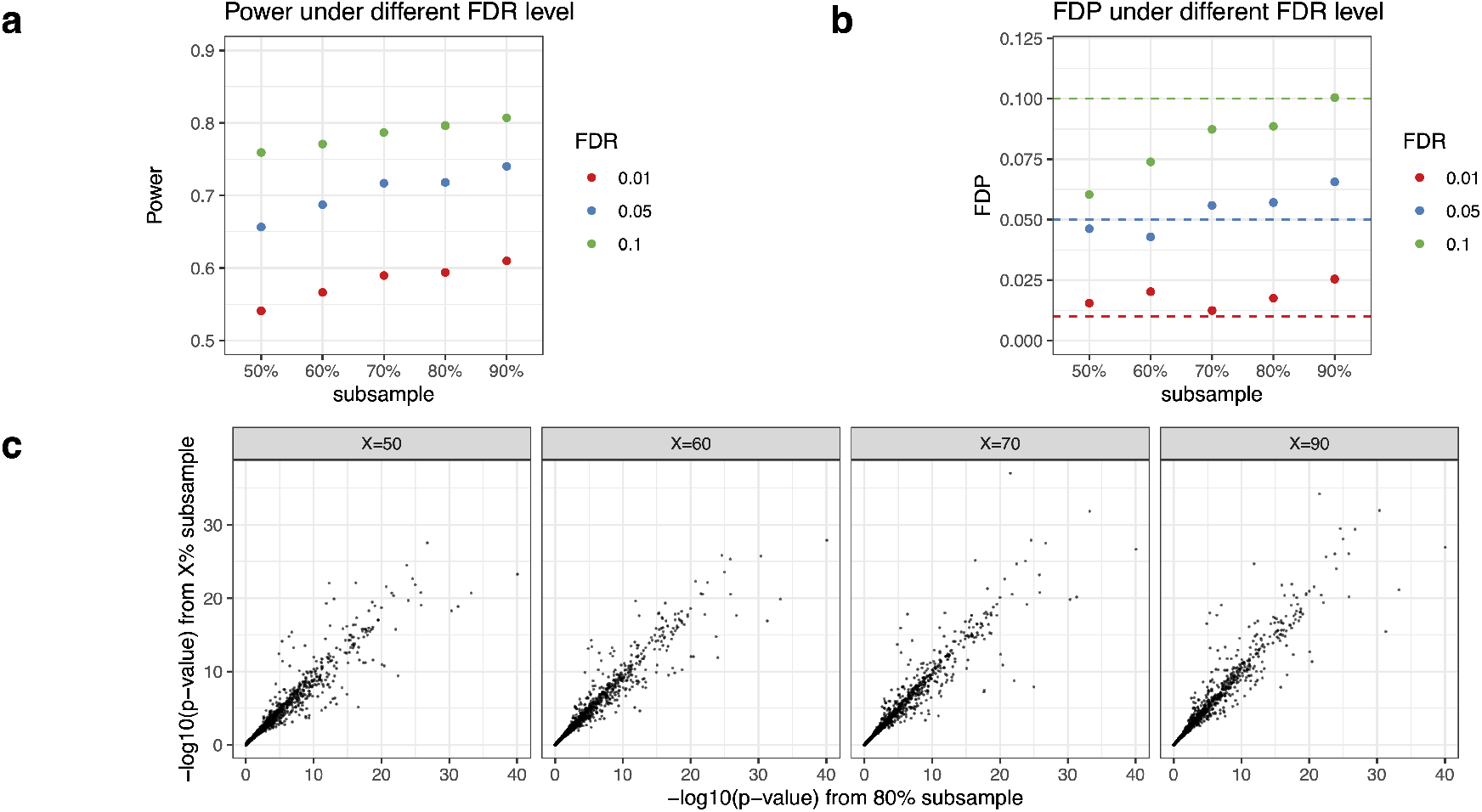
Robustness of PseudotimeDE to the subsampling proportion. Results are based on the synthetic high dispersion dataset and pseudotime inferred by Slingshot. **(a)** Power of PseudotimeDE using different subsampling proportions under FDR levels 0.01, 0.05, and 0.1. **(b)** FDP of PseudotimeDE using different subsampling proportions under FDR levels 0.01, 0.05, and 0.1. **(c)** Scatter plots of the default *p*-values using 80%as the subsampling proportion vs. the *p*-values using 50%, 60%, 70%or 90%as the subsampling proportion. The strong linearity (Pearson correlation coefficient *R* ≥ 0.96) of *p*-values under different subsampling proportions confirms the robustness of PseudotimeDE to the subsampling proportion.

## Notes

### Competing Interest Statement

The authors have declared no competing interest.

### Summary of Updates

We have corrected some typos in the text and figures.

